# Heterogeneous Receptor Expression Underlies Non-uniform Peptidergic Modulation of Olfaction in *Drosophila*

**DOI:** 10.1101/2022.04.27.489804

**Authors:** Tyler R. Sizemore, Julius Jonaitis, Andrew M. Dacks

**Affiliations:** Department of Biology, Life Sciences Building, West Virginia University, Morgantown, WV, 26506, USA; Department of Molecular, Cellular, and Developmental Biology, Yale Science Building, Yale University, New Haven, CT, 06520-8103, USA; Department of Neuroscience, West Virginia University, Morgantown, WV, 26506, USA

**Keywords:** Neuromodulation, Neuropeptide, Olfaction, Connectomics, Local Interneuron, *Drosophila*

## Abstract

Sensory systems are dynamically adjusted according to the animal’s ongoing needs by neuromodulators, such as neuropeptides. Although many neuropeptides are often widely-distributed throughout sensory networks, it is unclear whether such neuropeptides uniformly modulate network activity. Here, we leverage the numerically tractable primary olfactory center of *Drosophila* (the antennal lobe, AL) to resolve whether one such widely-distributed neuropeptide (myoinhibitory peptide, MIP) uniformly modulates AL processing. We find that despite being uniformly distributed across the AL, MIP decreases olfactory input to some glomeruli, while simultaneously increasing olfactory input to other glomeruli. We reveal that a heterogeneous ensemble of local interneurons (LNs) are the sole source of MIP within the AL. Through high-resolution connectomic analyses, as well as *in vivo* physiology, we find that the non-uniform effects of MIP are not likely due to MIPergic LN intrinsic properties (e.g., synaptic inputs/outputs, odor-evoked responses, etc.). Instead, we show that differential expression of the inhibitory MIP receptor (sex peptide receptor, SPR) across glomeruli allows MIP to act on distinct intraglomerular substrates, thus enabling differential modulation of olfactory input. Our findings demonstrate how even a seemingly simple case of modulation (i.e., a single neuropeptide acting through a single receptor) can have complex consequences on network processing by acting non-uniformly within different components of the overall network.

## Introduction

Animals use their sensory systems to internalize and process information about the identity, intensity, and valence of external stimuli, so they can properly navigate their environment. However, constant ecological and internal state fluctuations threaten the animal’s ability to accurately represent these stimuli. To address the demands these fluctuations impose, sensory systems use processes such as neuromodulation to flexibly adjust sensory processing and behavior. The largest and most ancient collection of neuromodulators are small peptides (~3-100 amino acids) termed neuropeptides^1–10^. For instance, neuropeptide F (NPF)/neuropeptide Y (NPY) play a conserved role in promoting feeding behaviors in sea slugs, humans, flies, zebrafish, nematodes, mosquitoes, and rodents^11–19^. Yet, despite their ubiquity and clear importance in nervous systems^20–22^, the mechanistic basis of peptidergic modulation of sensory processing remains unclear.

Often peptidergic modulation is strongly associated with a given physiological drive, and in some cases the actions of a neuropeptide within a single network can be associated with different behavioral contexts. For instance, myoinhibitory peptide (MIP) is necessary and sufficient to stimulate the drive of *Drosophila* towards food-odors within the context of satiation^23^, and is implicated in the post-mating shift in odor-preferences^24^. However, MIP appears to be uniformly distributed across all olfactory channels (“glomeruli”) that comprise the antennal lobe (AL), which processes far more than just food-related odors. Therefore, how can a ubiquitously distributed neuropeptide have stimulus-specific effects on sensory processing? Here, we reveal the cellular, physiological, and structural substrates that enable MIP to differentially modulate olfactory input to distinct olfactory channels.

## Results

### MIP differentially modulates olfactory input to distinct glomeruli

To test whether odorant responses within different glomeruli are uniformly modulated by MIP, we chose an odor which activates several glomeruli visible at the same/nearly the same imaging depth, and whose pattern of glomerular activation is well-established - apple cider vinegar (ACV)^25^. In this way, any non-uniform effects of MIP on OSN odor-responses across different glomeruli would be readily detectable. Furthermore, we recorded the odor-evoked responses of OSN axon terminals (i) before MIP was pressure injected onto the AL (“pre-MIP injection”), (ii) after MIP was injected, but before it was removed from the perfusate (“MIP”), and (iii) after a brief washout period (“post-washout”) **(see Methods) (Fig. 1a)**.

**Fig. 1.**
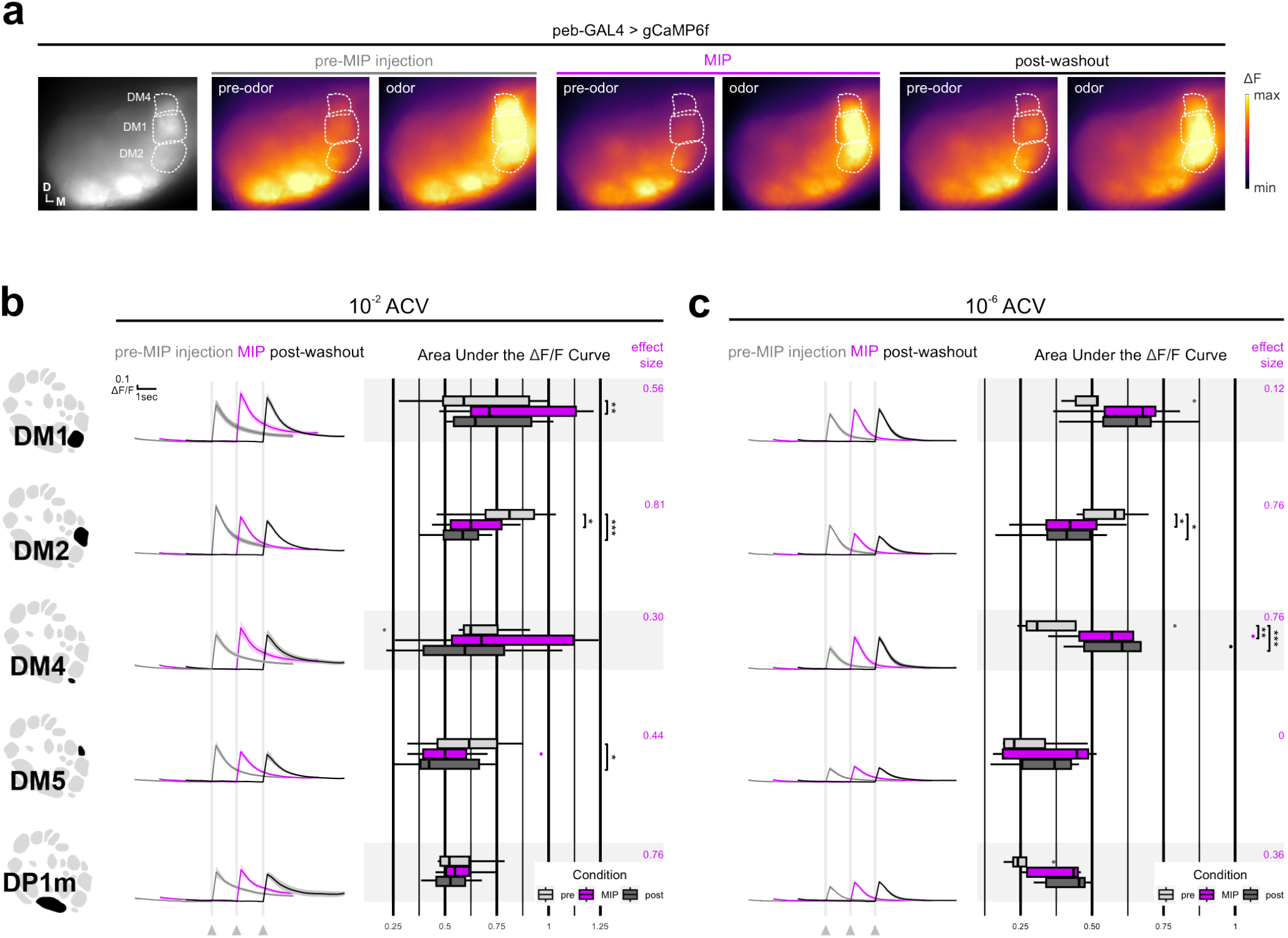
MIP differentially modulates OSN ACV responses. **(a)** Representative pseudocolored heatmaps of OSN GCaMP before and during odor presentation in several test glomeruli (dotted outlines). In each case, each odor presentation heatmap pair is grouped by stage of MIP pharmacological application (e.g., “pre-MIP injection). **(b)** DM1, DM2, DM4, DM5, and DP1m OSN responses to 10^-2^ ACV before (“pre-MIP injection”), after MIP pressure injection (“MIP”), and post-washout (“post-washout”). MIP significantly increases DM1 OSN responses (p = 0.004, pre-MIP injection vs. MIP AUC, n = 8; Holm-corrected RM t-tests). Conversely, MIP significantly decreases DM2 and DM5 OSN responses (<u>DM2</u>: p = 0.013, pre-MIP injection vs. MIP AUC & p = 0.001, pre-MIP injection vs. post-washout AUC, n = 8; Holm-corrected RM t-tests; <u>DM5</u>: p = 0.02, pre-MIP injection vs. post-washout, n = 8; Holm-corrected RM t-tests). In every case, effect size measurements are provided to the right of each set of AUC boxplots. **(c)** Same as **b**, but in response to 10^-6^ ACV. MIP significantly decreases DM2 OSN responses (p = 0.013, pre-MIP injection vs. MIP AUC & p = 0.01, pre-MIP injection vs. post-washout AUC, n = 5; Holm-corrected RM t-tests). Conversely, MIP significantly increases DM4 OSN responses (p = 0.002, pre-MIP injection vs. MIP AUC & p = 0.001, pre-MIP injection vs. post-washout AUC, n = 5; Holm-corrected RM t-tests). In every case, effect size measurements are provided to the right of each set of AUC boxplots. For each response: vertical & horizontal scale bars = 0.1 ΔF/F & one-second (respectively). Odor onset is indicated by vertical lines running up each column of traces. Statistical measures of effect size (either Kendall’s *W* or Cohen’s *d*) are provided to the right of each set of AUC boxplots. Glomerular schematics derived from an *in vivo* AL atlas^167^.

Before MIP application, OSNs robustly respond to both test concentrations of ACV **(Fig. 1b & 1c)**, then after MIP is pressure injected into the AL, DM1 OSN responses to 10^-2^ ACV are increased **(Fig. 1b)**. Similarly, DM4 OSN responses to 10^-6^ ACV are also increased after peptide application **(Fig. 1c)**. After a brief washout period, DM1 OSN responses to 10^-2^ ACV return to pre-peptide application responses **(Fig. 1b)**, whereas the increased DM4 OSN responses to 10^-6^ ACV are sustained **(Fig. 1c)**. In contrast, DM2 OSN responses are substantially diminished after peptide application regardless of odor concentration, and remain so post-washout **(Fig. 1b & 1c)**. Moreover, DM5 OSN responses to 10^-2^ ACV, which were decreased (albeit insignificantly) upon peptide application, become significantly diminished post-washout relative to pre-peptide application **(Fig. 1b)**.

Altogether, these results show that MIP can differentially modulate OSN odor-evoked responses in a glomerulus- and stimulus concentration-dependent manner, while also having concentration-independent consequences on OSNs of another glomerulus. However, these observations could be explained by differences in MIP-SPR signaling substrates across these glomeruli. For instance, there may be no synaptic input to DM1 OSNs from MIP-ir AL neurons, and therefore our prior observations **(Fig. 1b & 1c)** are the result of polysynaptic influences induced by MIP application. Therefore, we sought to test our suppositions by resolving the entire MIPergic signaling circuit architecture, including the identity of the presynaptic MIP-releasing AL neurons, their pre- & postsynaptic partners, and those postsynaptic partners that express SPR.

### Patchy GABAergic LNs are the sole source of MIP within the *Drosophila* AL

Previous neuroanatomical investigations suggested that the neurites of the AL-associated MIPergic neurons appear restricted to the AL, which implies MIP is released from AL LNs^26^. However, the *Drosophila*

AL houses ~200 LNs whose distinct roles in olfactory processing have been associated with their transmitter content and morphology^27–37^. For example, individual cholinergic AL LNs innervate many glomeruli and perform lateral excitation as a means for broadening odor representations in the AL^33,34,38–40^. Therefore, to resolve whether MIP-immunoreactive (MIP-ir) AL neurons are indeed AL LNs, and if they belong to a known AL LN chemical class, we assessed the overlap of MIP-immunoreactivity with markers for the major *Drosophila* small-neurotransmitters^41^ **(Fig. 2a and Supplementary Fig. 1)**. We find that AL MIP-ir neurons do not overlap with choline acetyltransferase (ChAT) or vesicular glutamate transporter (VGlut), but all MIP-ir neurons in the AL overlap with GAD1 (9.1±0.19 neurons, n = 5 brains, 10 ALs) **(Fig. 2a and Supplementary Fig. 1)**. In accordance with RNA-sequencing^42–44^, we find no detectable MIP-immunoreactive OSNs **(Supplementary Fig. 1)**. Altogether, these results suggest that an ensemble of ~9 GABAergic LNs are the source of MIP within the *Drosophila* AL.

**Fig. 2.**
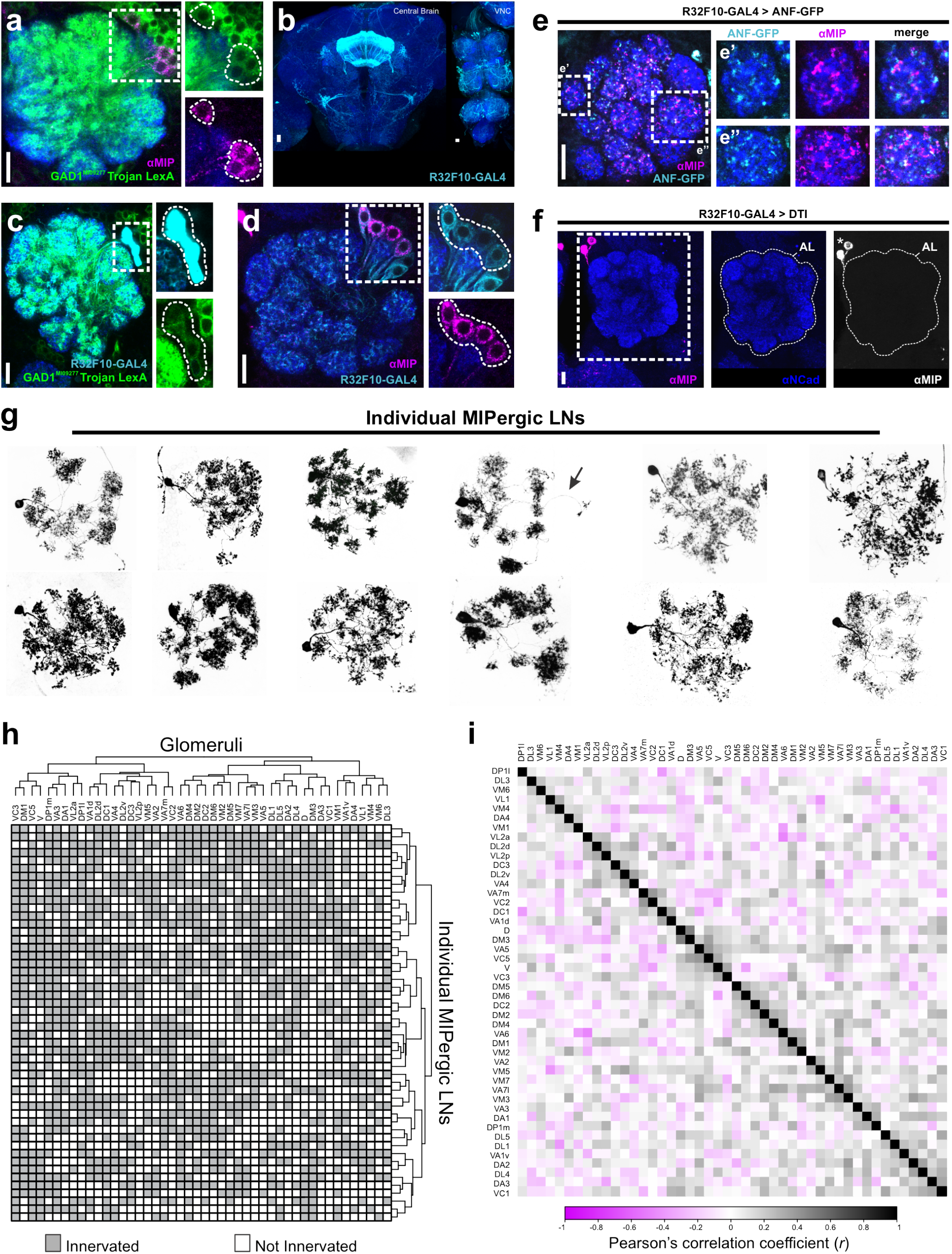
Myoinhibitory peptide (MIP) is released by GABAergic patchy LNs in the AL. **(a)** A protein-trap Trojan LexA driver for glutamic acid decarboxylase (GAD1), the rate-limiting enzyme for GABA, highlights all MIP-immunoreactive neurons in the AL. Cell counts: n = 5 brains, 10 ALs. **(b)** R32F10-GAL4 expression in the central brain and ventral nerve cord (VNC). **(c)** All R32F10-GAL4 AL LNs colocalize with the GAD1 Trojan LexA protein-trap driver. Cell counts: n = 5 brains, 9 ALs. **(d)** All MIP immunoreactive AL neurons (~8.7±0.3 neurons) are highlighted by R32F10-GAL4, which labels ~13.2 (±0.68) AL neurons in total. In addition to labeling all MIP-ir AL neurons, R32F10-GAL4 also labels ~4.5 (±0.68) non-MIPergic GABAergic AL neurons. Cell counts: n = 5 brains, 9 ALs. **(e)** Representative image of GFP-tagged rat preproatrial natriuretic factor (ANF-GFP) expression in R32F10-GAL4 AL LNs. Note the accumulation of ANF-GFP with MIP-ir punctae in R32F10-GAL4 AL LN terminals, such as those in two representative glomeruli DM5 **(e’)** and VA1d **(e”). (f)** Expression of a temperature-sensitive genetically encoded diphtheria toxin (DTI) in R32F10-GAL4 AL LNs abolishes all MIP-immunoreactivity (MIP-ir) in the AL. Note, because this ablation method is cell-specific, the median bundle cluster of MIP-ir neurons (the “MBDL” cluster; **see Supplementary Fig. 1**) outside of the AL remain intact when R32F10-GAL4 AL LNs are ablated (asterisks). **(g)** Stochastic labeling of individual R32F10-GAL4 AL LNs reveals MIP is released by patchy LNs. Arrow indicates a projection into the contralateral AL. **(h)** Glomerular innervation patterns of 50 individual MIPergic LNs organized by hierarchical clustering similarity. Each row represents the innervation pattern of a single clone, and each column represents a given glomerulus. Note that in some cases a clone might project into the contralateral AL, but here only the ipsilateral innervation patterns were included for analysis. **(i)** All pairwise correlations of MIPergic LN innervation patterns between AL glomeruli. Values correspond to the Pearson’s correlation coefficient for each glomerulus pair. In all cases: neuropil was delineated by anti-DN-Cadherin staining; scale bars = 10μm.

The adult *Drosophila* AL houses a variety of distinct GABAergic LNs, which can be subdivided into five major morphological types: panglomerular, multiglomerular, oligoglomerular, continuous, and patchy^32^. Like cortical interneurons^45,46^, these different interneuron morphological types play distinct roles in AL olfactory processing. To determine morphological type to which the MIPergic LNs belong, we screened the Janelia FlyLight driver line collection^47^, tested ~25 of those lines for MIP-immunoreactivity, and identified a GAL4 driver (R32F10-GAL4) that selectively highlights MIPergic LNs within the AL **(Fig. 2b-2d)**. We then combined R32F10-GAL4 with a GFP-tagged rat preproatrial natriuretic factor (ANF-GFP) which, when expressed within peptidergic neurons, is proteolytically processed and packaged into secretory vesicles and preferentially accumulates in peptidergic synaptic terminals^48^. We find broad ANF-GFP accumulation in R32F10-GAL4 AL LN terminals across the AL, confirming these LNs possess the necessary subcellular machinery for neuropeptide processing, packaging, and release **(Fig. 2e)**. Furthermore, all MIP-ir is abolished in the AL when R32F10-GAL4 AL LNs are ablated via temperature-gated expression of a cell-specific diphtheria toxin **(Fig. 2f)**. We then leveraged our selective genetic access to these LNs to resolve the morphology of individual MIPergic LNs through stochastic labeling^49^. From these experiments, we find that all MIPergic LNs have a discontinuous innervation pattern resembling that of patchy AL LNs **(Fig. 2g and Supplementary Fig. 1)**. Taken together, our data suggests that ~9 GABAergic patchy LNs are the sole source of MIP within the *Drosophila* AL.

There are many AL neurons, including other LNs, that are not patchy LNs but resemble the discontinuous morphology described for patchy LNs^32^. However, individual patchy LNs are unique in that they innervate different sets of glomeruli from animal-to-animal^32^. Therefore, we analyzed the set of glomeruli innervated by 50 individual MIPergic LNs and observed 50 distinct innervation patterns, thus demonstrating that no individual MIPergic LN innervates the same set of glomeruli across animals **(Fig. 2h and Supplementary Figs. 2 & 3)**. Additionally, we find individual MIPergic LNs do not preferentially innervate any one glomerulus over others **(Supplementary Fig. 2)**. When sister clones were assessed, we find that two individual MIPergic LNs co-innervate ~12 glomeruli (n = 5 brains, 5 sister clones) **(Supplementary Fig. 3a & 3b)**, and individual MIPergic LNs consistently innervated at least one of the hygro-/thermosensory associated glomeruli **(Supplementary Fig. 3c-3e)**. These results suggest that at least two MIPergic LNs innervate any single glomerulus, including hygro-/thermosensory glomeruli^50–52^. Moreover, these observations also demonstrate that individual MIPergic LNs innervate different glomeruli from animal-to-animal.

Most odorants activate more than one glomerulus in the *Drosophila* AL^25,53–55^. Thus, if individual MIPergic LNs innervate different sets of glomeruli from animal-to-animal, are there pairs of glomeruli that are innervated significantly more than others? If so, what ecological relationships exist amongst significantly correlated pairs of glomeruli? To determine the probability that an individual MIPergic LN that innervates one glomerulus will innervate/avoid another glomerulus, we leveraged our previous clonal analysis data **(Fig. 2g & 2h)** to calculate a correlation coefficient for all possible pairs of glomeruli **(Fig. 2i)**. This analysis revealed several statistically significant relationships, of which the most significant pairs were DM3-D (r = 0.49, p = 2.7 x 10 ^4^) and VL2p-VA6 (r = −0.47, p = 4.9 x 10^-4^) **(Fig. 2i and Supplementary Table 1)**. In addition to DM3-D and VL2p-VA6, this analysis also revealed a significant probability for MIPergic LN co-innervation amongst several pairs of glomeruli responsive to ACV^25^, such as VM2-DM1 (r = 0.35, p = 0.01), DM4-DM2 (r = 0.31, p = 0.03), and DP1m-DM1 (r = 0.29, p = 0.04). This suggests that the glomerular innervation patterns of individual MIPergic LN likely do not explain non-uniform MIPergic modulation of OSN **(Fig. 1)**.

However, it is plausible that differential modulation of OSN responses by MIP arises as a consequence of the non-uniform MIPergic LN pre-/postsynaptic sites across these glomeruli. Therefore, we sought to determine whether MIPergic LN input/output sites were heterogeneously distributed to any particular glomeruli throughout the AL.

### MIPergic LNs provide and receive broad input and output across the AL

We have shown that no individual MIPergic LN innervates the same set of glomeruli from animal-to-animal **(Fig. 3a)**, but every glomerulus is innervated by at least one MIPergic LN across all animals **(Fig. 3b)**. Therefore, we wondered whether significant differences in MIPergic LN input/output between glomeruli exist **(Fig. 3c)**, which would explain the non-uniform effects of MIPergic modulation. To test this, we measured the density of MIPergic LN-expressed mCD8::GFP, anti-MIP immunoreactive puncta, and the synaptic polarity markers DenMark and synaptotagmin.eGFP (syt.eGFP)^56,57^ in each glomerulus across many animals **(Fig. 3d & 3e)**. We find that the density of each indicator varies across glomeruli but are stereotypic across samples **(Fig. 3e and Supplementary Fig. 3f-3i)**. The density of the output indicators (syt.eGFP and MIP-ir puncta) were statistically correlated, and nearly every indicator scaled with MIPergic LN intraglomerular cable density **(Supplementary Fig. 3f-3i)**. Even so, we find within-indicator voxel densities are generally evenly distributed across each glomerulus, suggesting MIPergic LN input and output are evenly distributed across the AL **(Fig. 3e)**.

**Fig. 3.**
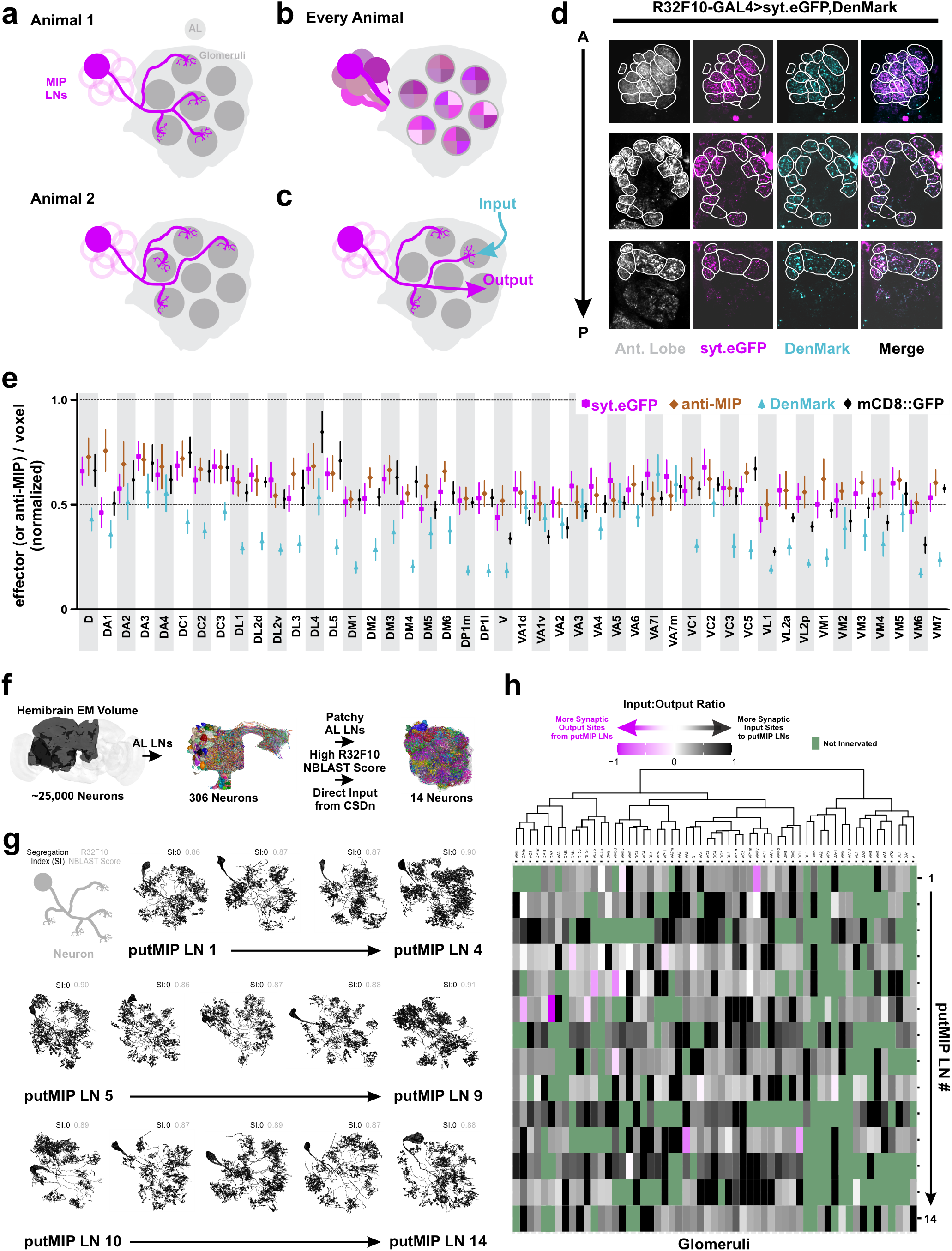
MIPergic LN input and output site throughout the entire AL. **(a)** Individual MIPergic LNs project to different glomeruli from animal-to-animal. **(b)** The MIPergic LN ensemble covers the entire AL across all animals. **(c)** Do MIPergic LNs receive input from particular sets of glomeruli? Are there particular sets of glomeruli subject to more/less MIPergic LN output than others? **(d)** Representative image of glomerular voxel density analysis. Here, MIPergic LNs express synaptotagmin.eGFP (syt.eGFP; magenta) and DenMark (cyan) and their respective density is measured within each AL glomerulus (Ant. Lobe; grey). Glomeruli outlined in white. **(e)** syt.eGFP (magenta), DenMark (cyan), mCD8::GFP (black) and anti-MIP (brown) puncta density per voxel within each AL glomerulus. Each indicator is normalized to the highest value within that indicator. Data are represented as the mean±SEM of each indicator’s voxel density within a given glomerulus. For each indicator, n = 7 (syt.eGFP), 7 (DenMark), 4 (mCD8::GFP), 4 (anti-MIP) brains. **(f)** Schematic representation of procedures used to identify ideal putative MIPergic LN (putMIP LN) candidates from the FlyEM FIB-SEM hemibrain connectome volume **(see Methods). (g)** putMIP LN mesh skeletons identified from the hemibrain EM volume. For each neuron, values in the upper right-hand corner are that neuron’s synaptic segregation index (black) and GMR32F10-GAL4 NBLAST similarity score (grey). **(h)** putMIP LN intraglomerular input:output ratio across the AL. Each column represents a given glomerulus, and each row represents the input:output ratio of a single putMIP LN. Glomeruli not innervated by the given putMIP LN are green. Glomeruli are organized by hierarchical clustering similarity. Data only consider putMIP LN connections within the ipsilateral AL.

These puncta analyses afford the advantages of analyzing MIPergic LN synaptic polarity across many individuals of both sexes. However, light microscopy is limited by its inability to resolve fine structures such as axons/dendrites^58,59^. Therefore, we performed similar analyses on individual putative MIPergic LNs (putMIP LNs) within the most densely-reconstructed *Drosophila* central brain EM volume to-date, the hemibrain^60,61^. Additionally, EM-level analyses of putMIP LN connectivity have the added benefit of shedding light on what type(s) of neuron(s) and/or stimuli might generally promote MIP recruitment in AL processing. More specifically, EM analysis of putMIP LN connectivity allowed us to determine: (*i*) do certain glomeruli receive more input from putMIP LNs (and vice versa) than others? (*ii*) what neurons are upstream/downstream of putMIP LNs in each glomerulus? and, (iii) at which putMIP LN presynaptic terminals are vesicles associated with neuropeptides (dense core vesicles, DCVs)^62,63^ found? Thus, we first used several criteria **(see Methods)** to identify fully-reconstructed putMIP LNs, which resulted in the identification of 14 ideal candidates **(Fig. 3f and Supplementary Table 1)**.

After identifying several optimal candidates, we tested whether any putMIP LNs have distinct dendritic and axonic compartments. If true, this would suggest putMIP LNs make region-specific input/output, as has been suggested for the “heterogeneous LNs” in the honeybee AL^64–67^. Synaptic flow centrality and axonal-dendritic segregation indices^68^ reveal all putMIP LNs lack clearly separable input/output compartments **(Fig. 3g and Supplementary Table 1)**. When we assess the ratio of input-to-output along a given putMIP LN’s intraglomerular neurites, we find that the amount of input a given putMIP LN receives typically outnumbers the amount of putMIP LN output within any given glomerulus **(Fig. 3h)**. To better understand how these inputs might drive MIPergic modulation, we assessed the general identity of all inputs a putMIP LN receives, as well as the class and transmitter type of each presynaptic input an intraglomerular putMIP LN arbor receives.

Generally, nearly half of all putMIP LNs receive more input from other LNs than other principal neuron categories (6/14 putMIP LNs; ~38-40% total input) **(Fig. 4a)**. Similarly, just as many putMIP LNs receive the majority of their input from PNs than any other principal neuron category (6/14 putMIP LNs; ~31-33% total input) **(Fig. 4a)**. Additionally, we found that within a given glomerulus putMIP LNs largely avoid one another, but do occasionally form synaptic connections **(Fig. 4b and Supplementary Fig. 3j & 3k)**. When we refine these analyses by considering the putMIP LN’s intraglomerular connectivity and the presynaptic partner’s identity, we find the amount of excitatory or inhibitory input a given putMIP LN receives varies greatly across glomeruli. However, in every case, putMIP LNs generally receive far more excitatory than inhibitory input within any given glomerulus (~65-93% of all glomeruli innervated by the given putMIP LN) **(Fig. 4b)**. This suggests that MIPergic LNs intraglomerular processes may be broadly activated by disparate odorants, which would suggest uniform release of MIP may occur in response to various odors. Therefore, we tested whether MIPergic LNs are broadly activated *in vivo* by chemically diverse odorants.

**Fig. 4.**
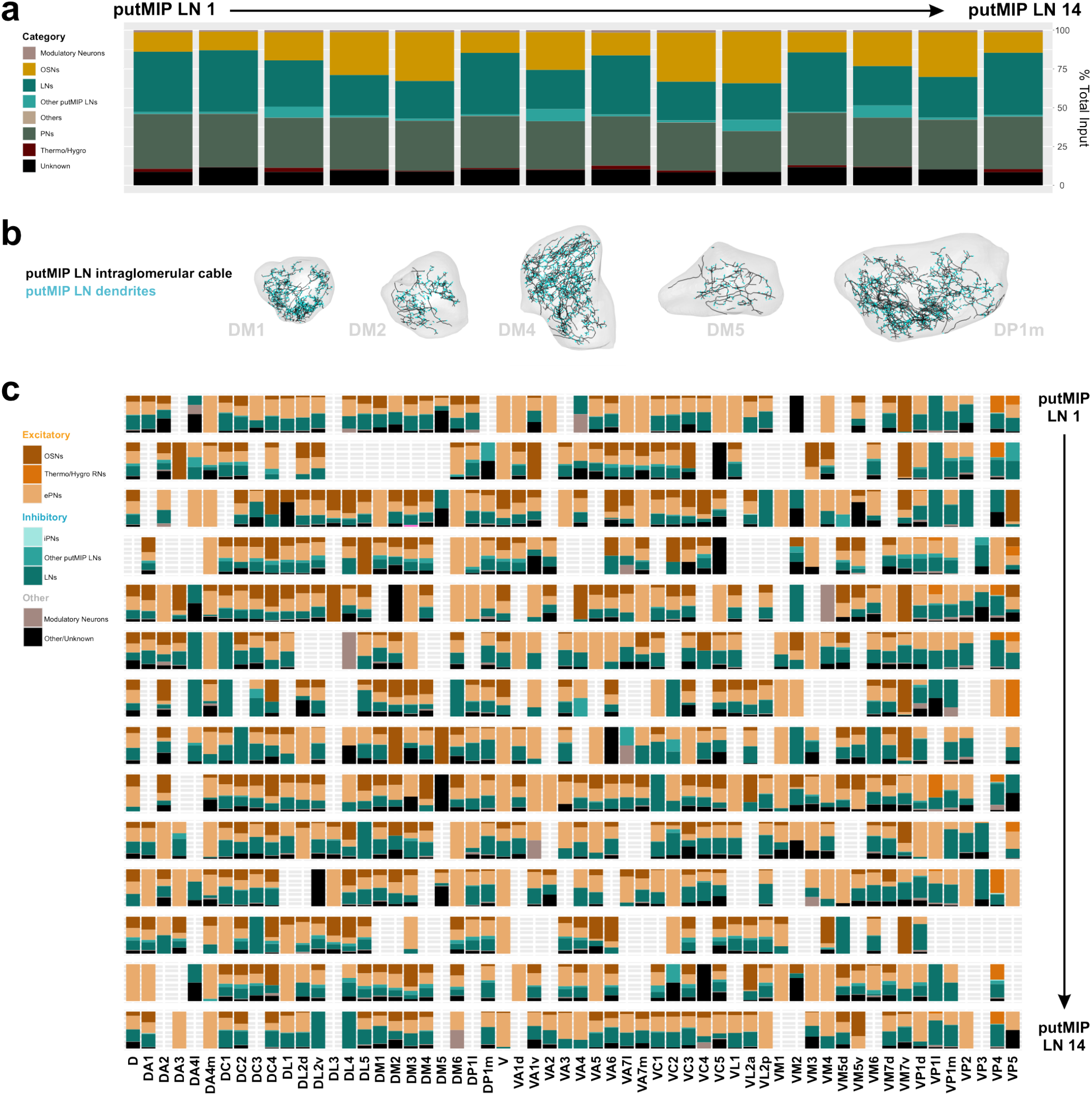
Anatomical inputs to putMIP LNs and functional glomerular outputs from identified MIPergic LNs. **(a)** putMIP LN upstream partners’ demographics. Data are represented as a function of the total amount of input a putMIP LN receives from all categories. **(b)** Representative images of putMIP LN intraglomerular cable and dendrites plotted within their corresponding glomeruli. **(c)** The amount of excitatory, inhibitory, and modulatory input each putMIP LN receives within every glomerulus, broken apart by presynaptic neuron identity, and represented as a function of the total amount of input a given putMIP LN receives within the glomerulus. Glomeruli with no bar graph are those that the given putMIP LN does not innervate.

### MIPergic LNs generally display glomerulus-specific odor-evoked responses

Synapse counts have been shown to strongly predict functional output strength in neurons within other systems, including other *Drosophila* AL neurons^28,69–74^. Our connectomic analyses intraglomerular putMIP LN arbors generally receive mostly excitatory input within a given glomerulus **(Fig. 4b)**, which would imply that MIPergic LNs are broadly activated regardless of odor identity. This would be consistent with previous characterizations of large ensembles of GABAergic AL LNs odor-evoked GCaMP responses that have observed odor-invariant activation across nearly all glomeruli^54,75^. Therefore, we recorded the *in vivo* odor-evoked responses of MIPergic LN intra-glomerular neurites to a panel of chemically diverse odorants across multiple glomeruli **(Fig. 5a)**. Moreover, we chose to image within glomeruli whose cognate PNs respond to at least one odorant in our test panel to better understand what role the MIPergic LNs may play in the OSN-to-PN information transfer. For example, benzaldehyde and geranyl acetate each evoke responses in DM1, DM4, DP1l, VA2, and VM2 PNs^37,76–78^.

**Fig. 5.**
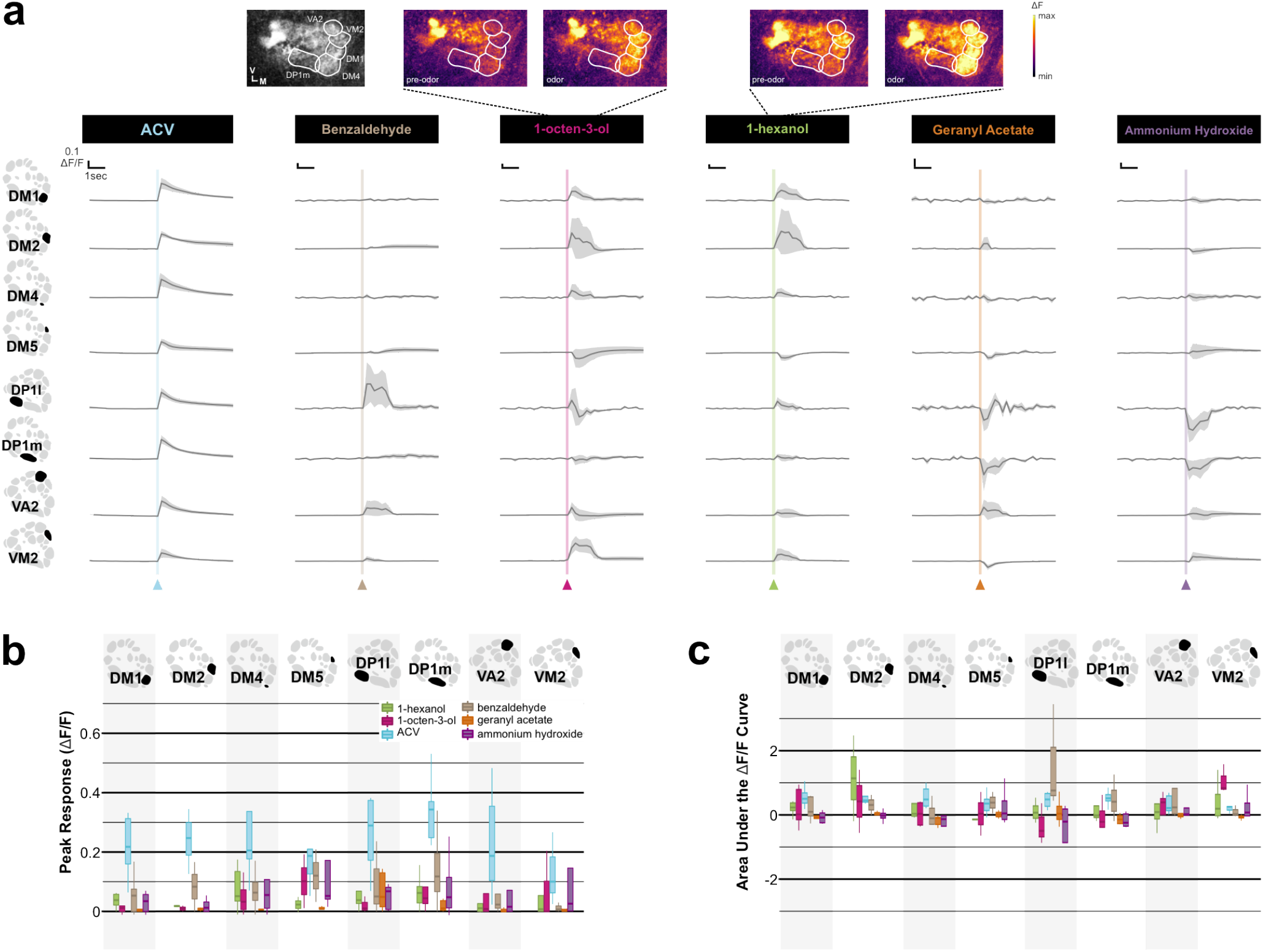
MIPergic LNs are differentially activated by different odors. **(a)** Odor-evoked responses of MIPergic LN neurites within several AL glomeruli (far left column). Odors tested were presented at 10^-2^ and include: apple cider vinegar (ACV), benzaldehyde, 1-octen-3-ol, 1-hexanol, geranyl acetate, and ammonium hydroxide. For each stimulus: n = 3-10 animals; vertical and horizontal scale bars = 0.1 ΔF/F & 1 second (respectively). Odor onset is indicated by the vertical lines running up each column of traces. **(b)** Peak response (ΔF/F) of MIPergic LN intraglomerular neurites from odor onset to ~1 second after stimulus onset across all glomeruli tested for each stimulus. **(c)** Area under the ΔF/F curve (AUC) of MIPergic LN intraglomerular neurites across glomeruli for each stimulus. In all cases: Glomerular schematics derived from an *in vivo* AL atlas^167^.

In contrast to other GABAergic AL LNs^54,75^, we find MIPergic LNs generally display glomerulus-specific responses to all test odors **(Fig. 5a)**. For example, MIPergic LN neurites within VM2 and DP1m – two glomeruli visible at the same imaging depth - are simultaneously activated and inhibited by 1-octen-3-ol, respectively **(Fig. 5a-5c)**. This same odor drove post-excitatory depression in MIPergic LN neurites within DP1l and VA2 **(Fig. 5a-5c)**. However, in many cases MIPergic LN intraglomerular neurites did not respond to the given odorant **(Fig. 5a-5c)**. This shows that, unlike other GABAergic AL LNs which are broadly activated in response to similar stimuli^33,54,75^, MIPergic LN intraglomerular processes are differentially activated by different odors. However, we acknowledge that these odor-evoked responses do not necessarily reflect MIP release itself, which we were unable to test for reasons described below **(see Discussion)**.

Notably, ACV elicited robust activation of MIPergic LN intraglomerular processes across all identifiable glomeruli **(Fig. 5a-5c)**, including those glomeruli we found MIP non-uniformly modulates **(Fig. 1)**. This finding, together with our earlier results **(Figs. 2-5)**, suggests that the non-uniform effects of MIP on olfactory input likely do not arise from the presynaptic MIP-releasing neurons themselves. Instead, these results suggest the non-uniform effects of MIP on olfactory input are an emergent property of either (*i*) MIPergic LN postsynaptic targets, and/or (*ii*) differential SPR expression across the AL.

### MIPergic LN downstream partners and widespread SPR expression within the AL

To determine the AL principal neurons likely targeted by MIPergic LNs, we first assessed the general output demographics for each putMIP LN **(Fig. 6a & 6b)**. Of the AL principal neuron types, most putMIP LNs chiefly target PNs (71% of putMIP LNs; ~19-31% of putMIP LN total output) **(Fig. 6a)**. The remaining minority of putMIP LNs are chiefly presynaptic to OSNs (29% of putMIP LNs; ~27-33% of putMIP LN total output) **(Fig. 6a)**. Since AL PNs express GABA_A_ and GABAB receptors^27^, these results would suggest that MIPergic LNs may provide fast- and slow-acting inhibition across the AL, perhaps as a means to normalize PN odor-evoked responses. To determine which of these downstream partners **(Fig. 6b)** are likely targeted by MIPergic modulation, we determined which post-synaptic partners were downstream of putMIP LN terminals where DCVs are observable **(Fig. 6c)**. We observed several instances where DCVs could be found in putMIP LN terminals presynaptic to OSNs, PNs, and ventral LNs **(Fig. 6c)**. However, MIPergic LNs could also release other neuropeptides, so the presence of DCVs in MIPergic LN presynaptic terminals does not necessarily mean the downstream neuron is modulated by MIP. Moreover, putMIP LN EM analyses indicate several AL principal neuron types are plausible targets for MIPergic modulation **(Figs. 3, 6, and 7a)**. To determine which downstream partners are subject to MIPergic modulation, we identified the AL neurons that express MIP’s cognate receptor, the inhibitory SPR receptor^79–82^. To do so, we used a CRISPR/Cas9-mediated T2A-GAL4 insertion within the endogenous SPR locus to enable GAL4 expression within SPR-expressing cells^83^ **(Fig. 7b)**.

**Fig. 6.**
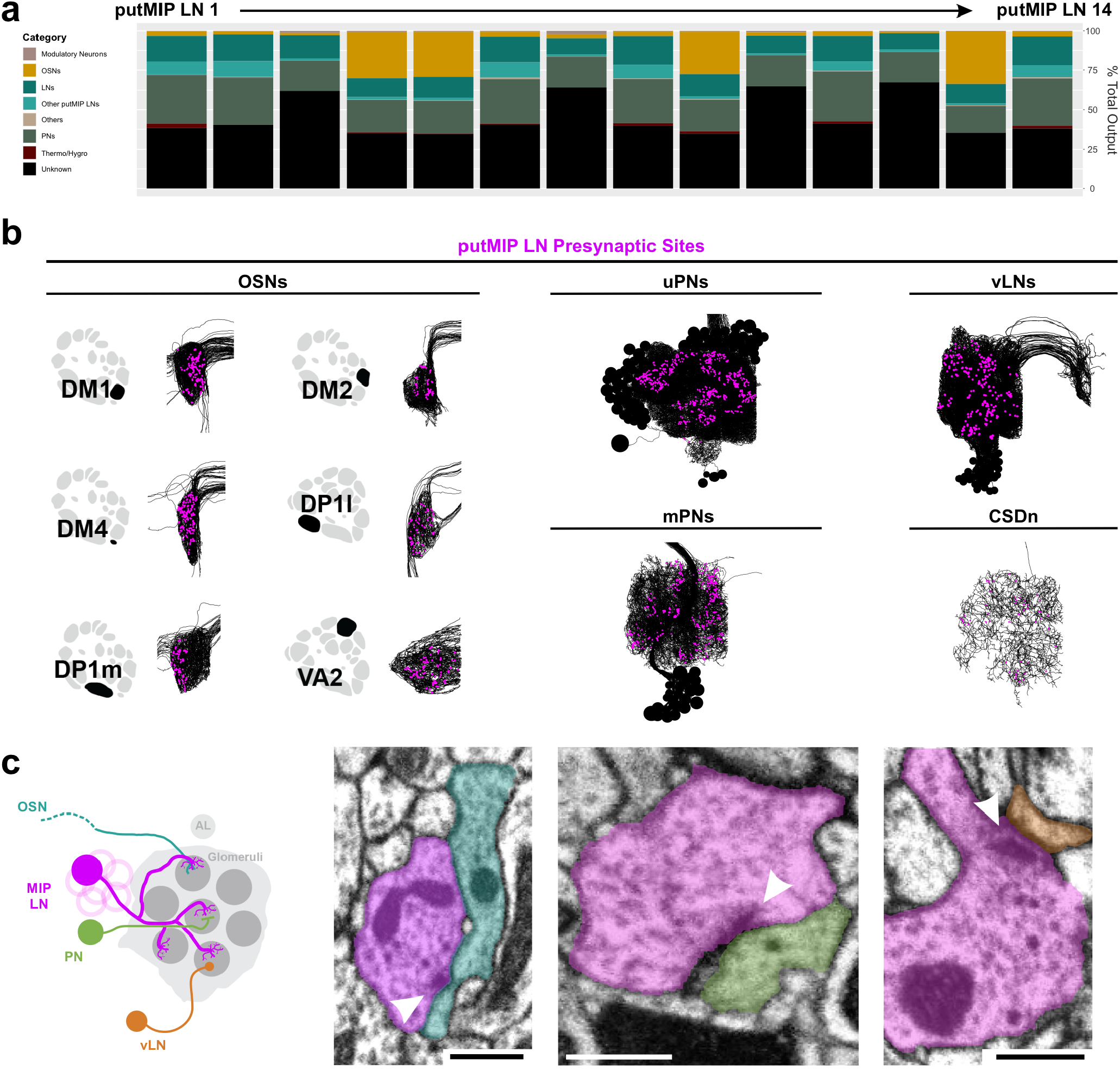
Postsynaptic targets of each putMIP LN and representative putMIP LN presynaptic terminals with dense core vesicles (DCVs). **(a)** Demographics of all putMIP LN postsynaptic targets by neuron type. Data are represented as a function of the total amount of output a putMIP LN sends to all categories. **(b)** Representative putMIP LN postsynaptic partner skeletons (black) with their respective putMIP LN presynaptic locations (magenta). Glomerular schematics derived from an *in vivo* AL atlas^167^. **(c)** Representative instances where DCVs in the putMIP LN presynaptic terminal. From left to right: DCVs are in putMIP LN presynaptic terminals upstream of OSNs (cyan), PNs (green), and ventral LNs (vLN; orange). In all cases: white arrowheads indicate the putMIP LN’s presynaptic site; scale bars = 500nm.

**Fig. 7.**
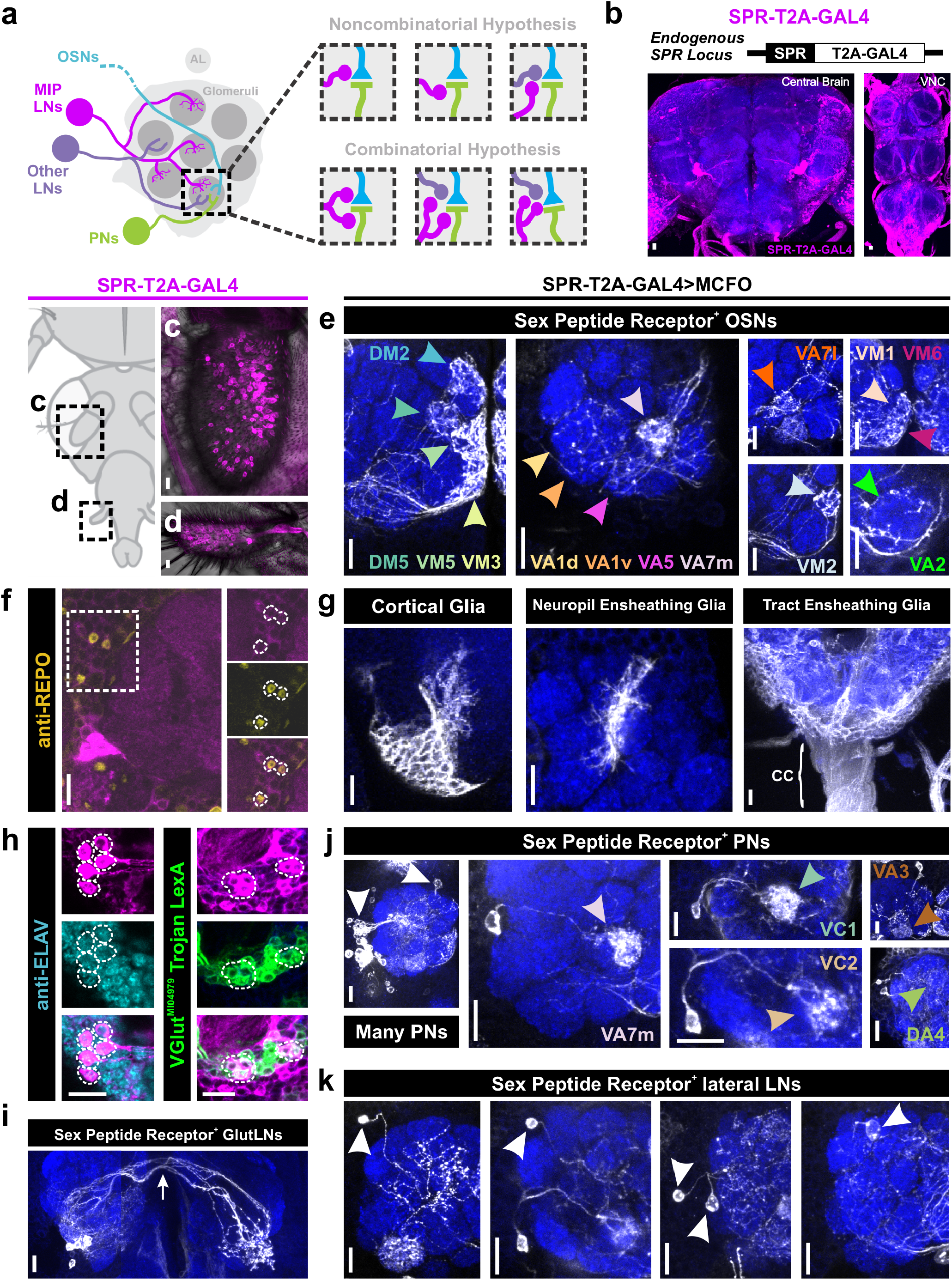
Widespread sex peptide receptor (SPR) expression throughout the AL. **(a)** MIPergic LNs (magenta) form synaptic connections with all principal neuron types in the AL; OSNs (cyan), PNs (green), and other LNs (purple). Therefore, within a single glomerulus, MIPergic modulation might target any one of these neuron types (“Non-combinatorial Hypothesis”), or multiple neuron types (“Combinatorial Hypothesis”). **(b)** SPR expression (magenta) revealed through a CRISPR/Cas9 T2A-GAL4 insertion in the SPR-coding intron. **(c & d)** SPR-T2A-GAL4 expression in OSNs in the third-antennal segment and maxillary palp. **(e)** SPR-T2A-GAL4 stochastic labeling experiments where the antennal nerve remains intact reveals SPR-expressing OSNs project to: DM2, DM5, VM5v, VM5d, VM3, VA1d, VA1v, VA5, VA7m, VA7l, VM1, VM6, VM2, and VA2. **(f)** SPR-T2A-GAL4 colocalizes with the glial marker REPO (yellow). **(g)** SPR-T2A-GAL4 stochastic labeling reveals expression in cortical, neuropil ensheathing, and tract ensheathing glia. “CC” = Cervical Connective. **(h)** Several SPR-T2A-GAL4 neurons are immunopositive for the proneural marker ELAV (cyan), a subset of which colocalize with VGlut^MI04979^ Trojan LexA (green). **(i)** SPR-T2A-GAL4 stochastic labeling reveals several bilaterally-projecting ventral glutamatergic LNs (GlutLNs). White arrow = bilateral projection. **(j)** Several lateral and anterodorsal PNs (white arrowheads) are highlighted via SPR-T2A-GAL4 stochastic labeling, some of which project to: VA7l, VC1, VC2, VA3, and DA4. **(k)** Approximately five lateral LNs are identified through SPR-T2A-GAL4 stochastic labeling. In all cases: neuropil was delineated by anti-DN-Cadherin staining; scale bars = 10μm.

In *Drosophila,* OSN somata are located within the third-antennal segment and maxillary palp^84,85^. We find 208.9±11.89 (n = 17 animals, 30 antennae) and 63.42±4.31 (n = 18 animals, 31 maxillary palps) SPR-T2A-GAL4^+^ neurons in the third-antennal segment and the maxillary palp, respectively **(Fig. 7c & 7d)**. As there are ~945 and ~113 OSNs in the antennae and maxillary palps, respectively^86^, this would suggest that ~22% of antennal OSNs and ~56% of maxillary palp OSNs express SPR. The number of SPR-T2A-GAL4^+^ neurons in either appendage do not significantly differ based on the animal’s sex or mating status (<u>antennae</u>: p = 0.107, one-way ANOVA; <u>maxillary palps</u>: p = 0.559, Kruskal-Wallis test). However, this does not discount differences in the level of SPR expression within these neurons based on the animal’s sex or mating status. Through stochastic labeling experiments where the antennal nerve is left attached to the brain, we found OSN fibers that innervate many distinct glomeruli, including several ACV-responsive OSNs **(Fig. 7e and Supplementary Table 1)**. Interestingly, we found SPR-T2A-GAL4 expression in afferents belonging to every sensory modality **(Supplementary Fig. 4)**, which suggests MIPergic modulation of sensory afferents may be a fundamental feature in *Drosophila*.

Within the brain, we noted overlap between SPR-T2A-GAL4 and the glial marker reverse polarity (REPO) **(Fig. 7f)**, which we found correspond to: (1) cortical glia, (2) neuropil ensheathing glia, and (3) tract ensheathing glia **(Fig. 7g)**. However, there is no evidence directly linking the actions of these glial subtypes with AL processing^87–89^, so we turned our attention to SPR-T2A-GAL4 cells immunopositive for the proneural gene embryonic lethal abnormal vision (ELAV) **(Fig. 7h)**. Through intersectional genetics and stochastic labeling, we find that these neurons consist of: 4.89±0.21 (n = 23 brains, 44 ALs) SPR-expressing ventral glutamatergic LNs (GlutLNs) **(Fig. 7h & 7i)**, uniglomerular PNs **(Fig. 7j and Supplementary Table 1)**, and several lateral LNs **(Fig. 7k)**. In agreement with these results, we find similar neuron types using another SPR driver (SPR-GAL4::VP16)^90^ **(Supplementary Fig. 5)**, several publicly available scRNA-seq datasets^91–93^ **(Supplementary Fig. 6)**, and a novel SPR^MI13553^-T2A-LexA::QFAD driver **(Supplementary Fig. 7)**.

### Differential SPR expression across glomeruli enables non-uniform MIPergic modulation of olfactory input

To test the necessity of direct MIP-SPR signaling on modulation of OSN odor-evoked responses, we repeated our earlier experiments **(see Fig. 1)**, but used RNA interference (RNAi) to knockdown SPR specifically within OSNs **(Fig. 8a & 8b)**. Moreover, this SPR-RNAi has been used to effectively knockdown SPR in OSNs previously^24^, and abolishes SPR immunoreactivity in the *Drosophila* CNS when expressed pan-neuronally^94^. We find that SPR knockdown abolishes the MIP-induced decrease in the odor-evoked responses of DM2 and DM5 OSNs **(Fig. 8c & 8d)**. This result is consistent with SPR-expression in DM2 and DM5 OSNs **(Fig. 7e)**, and suggests MIP directly decreases the odor-evoked responses of DM2 and DM5 OSNs. In contrast, SPR knockdown in DM1 and DM4 OSNs does not prevent their responses from increasing after peptide application **(Fig. 8c & 8d)**. Since we did not observe SPR-expression in DM1 and DM4 OSNs **(Fig. 7)**, and SPR knockdown in these OSNs does not abolish the MIP-induced increase in their responses **(Fig. 8c & 8d)**, our results suggest MIP acts polysynaptically to disinhibit (thus, increasing) DM1 and DM4 OSN odor-evoked responses **(Fig. 8e)**.

**Fig. 8.**
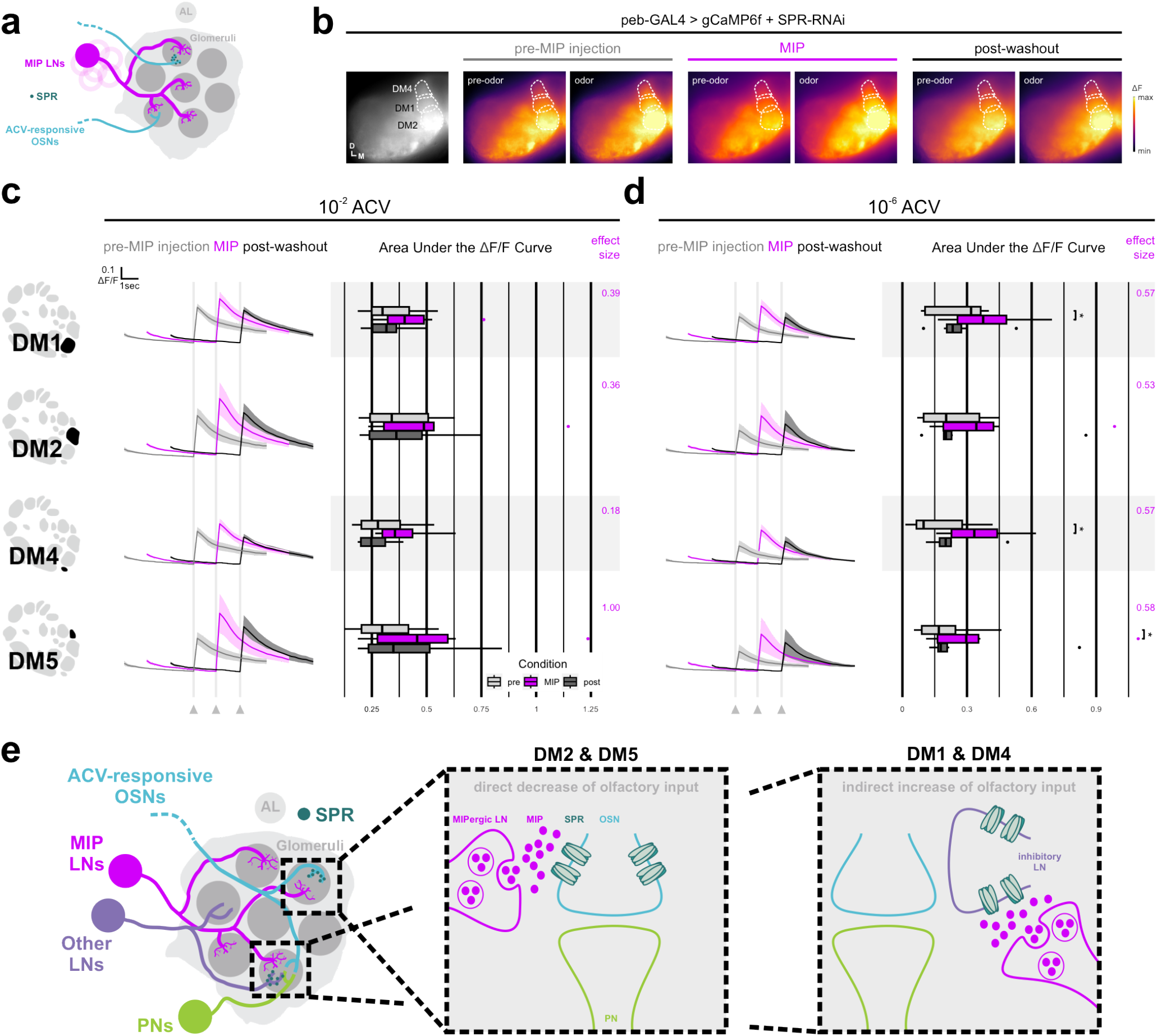
SPR knockdown in OSNs reveals heterogeneous SPR expression across glomeruli enables non-uniform MIPergic modulation of OSN ACV responses. **(a)** Individual MIPergic LNs (magenta) significantly co-innervate pairs of ACV-responsive glomeruli (cyan). Moreover, ACV-responsive OSNs (cyan) form synaptic connections with MIPergic LNs (magenta) and express the MIP receptor, SPR (turquoise). **(b)** Representative pseudocolored heatmaps of OSN GCaMP before and during odor presentation in several test glomeruli (dotted outlines) of animals where SPR is knocked down. In each case, each odor presentation heatmap pair is grouped by stage of MIP pharmacological application (e.g., “pre-MIP injection”). **(c & d)** SPR knockdown in OSNs abolishes MIP-induced decrease in DM2 and DM5 OSN responses (<u>DM2</u>: 10^-2^: p = 0.136, RM oneway ANOVA, n = 6; 10^-6^: p = 0.063, pre-MIP injection vs. MIP AUC & p = 0.688, pre-MIP injection vs. MIP AUC, n = 6; Holm-adjusted Wilcoxon signed-rank test; <u>DM5</u>: 10^-2^: p = 0.135, RM one-way ANOVA, n = 6; 10^-6^: p = 0.063, pre-MIP injection vs. MIP AUC & p = 0.313, pre-MIP injection vs. MIP AUC, n = 6; Holm-adjusted Wilcoxon signed-rank test). In contrast, SPR knockdown in OSNs does not abolish MIP-induced increases in DM1 and DM4 OSN responses (<u>DM1</u>: p = 0.031, pre-MIP injection vs. MIP AUC, n = 7; Holm-adjusted Wilcoxon signed-rank test; <u>DM4</u>: p = 0.031, pre-MIP injection vs. MIP AUC, n = 7; Holm-adjusted Wilcoxon signed-rank test). In every case, effect size measurements are provided to the right of each set of AUC boxplots. **(e)** Conceptual model of differential MIPergic modulation of OSN responses across multiple AL glomeruli. Our data show that the MIPergic LNs are the sole source of MIP to the AL, where MIP acts to directly decrease DM2 and DM5 OSN responses. Our data also show that MIP acts to indirectly increase DM1 and DM4 OSN responses, likely through disinhibition. For each response: vertical and horizontal scale bars = 0.1 ΔF/F & one-second (respectively). Odor onset is indicated by vertical lines running up each column of traces. Statistical measures of effect size (either Kendall’s *W* or Cohen’s d) are provided to the right of each set of AUC boxplots. Glomerular schematics derived from an *in vivo* AL atlas^167^.

## Discussion

### Seemingly simplistic circuitry gives rise to complex modulation

Our data reveals the circuit topology that enables a single neuropeptide, acting through a single receptor, to differentially modulate olfactory processing. We show that pharmacological application of MIP elicits non-uniform and complex effects on olfactory input to the *Drosophila* primary olfactory center. Here, MIP reduces the responses of OSNs in some glomeruli, and simultaneously enhances the responses of OSNs in other glomeruli **(Fig. 1)**. We show that the non-uniform effects of MIP on olfactory input is likely not an emergent property of the identity, structure, and/or connectivity of the MIP-releasing neurons, themselves. Instead, we find that differential SPR expression within distinct glomeruli enables MIP to non-uniformly modulate olfactory input across olfactory channels.

### Non-stereotypical neurons in a stereotyped neural network

We found that individual MIPergic LNs innervate a different repertoire of glomeruli across animals and do not preferentially innervate any one glomerulus over others **(Fig. 2 and Supplementary Fig. 2)**. These findings are consistent with earlier reports wherein patchy AL LNs were first generally described^32^. But, what factor(s) give rise to the tremendous flexibility within this single morphological subtype? One explanation might be that MIPergic LN morphological idiosyncrasy is a byproduct of experience during development. However, OSN removal in the adult does not disrupt the animal-to-animal variability of patchy LNs^32^. To the best of our knowledge, a single locus (e.g., environmental experience or heritable trait) that would support animal-to-animal variation in patchy LNs has not been identified.

Another explanation for animal-to-animal differences in individual MIPergic LN morphology is that it may not matter which individual MIPergic LN forms synapses with which downstream target, as long as all of the MIPergic LN downstream targets are met. Every nervous system is the byproduct of the adaptive pressures demanded by the animal’s niche; a place that can continually change in seemingly unpredictable ways. Therefore, a developmental “parameter space” may exist, wherein just enough genetic idiosyncrasy is allowed for to help prevent extinction in the face of environmental perturbations. The breadth of this developmental parameter space (or the degree of variability from the “median”) would be defined by many generations of selective pressures, wherein subtle changes in genetic idiosyncrasies might equally result in winners and losers. As a consequence of these genetic idiosyncrasies, phenotypic variability in a given developmental program would inevitably accumulate, resulting in the observed animal-to-animal variability in neuronal features (e.g., morphology, ion channel distribution, etc.). Consistent with this idea, animal-to-animal variations in neural circuitry have been noted in grasshoppers^95^, crabs^96–100^, lobsters^101,102^, flies^32,103,104^, and rats^105^. Moreover, inter-animal variations in neuronal architecture are one of several features implicated in interanimal behavioral variations^104,106–110^. However, despite this variability, overall neuronal circuit functions persist including consistent MIPergic LN synaptic polarity marker density **(Fig. 3)**, MIPergic LN within-odor responses **(Fig. 5)**, MIP-induced decreases in DM2 and DM5 OSN responses across animals **(Fig. 6)**, and SPR expression **(Fig. 7)**. Moreover, several positive and negative correlations exist for pairs of glomeruli innervated by single MIPergic LNs, such as the significant probability for MIPergic LN co-innervation in ACV-responsive glomeruli **(Fig. 2i and Supplementary Table 1)**. Together, these results suggest that the morphology of an individual MIPergic LNs can differ from animal-to-animal, as long as the right combinations of downstream targets (e.g., ACV-responsive neurons) are met by the ensemble.

### Functional implications of GABA and MIP co-transmission from MIPergic AL LNs

We have shown that a small ensemble (~5% of all AL LNs) of GABAergic AL LNs are the sole source of MIP in the *Drosophila* AL **(Fig. 2 and Supplementary Fig. 1)**. This implies that MIPergic LNs have the capacity to adjust AL olfactory processing through both GABA- and MIP-release. Previous works found that the iono- and metabotropic GABA receptors are expressed amongst all AL principal neuron types^27,29,111^, and we show SPR is analogously expressed by members of every AL principal neuron type **(Fig.7)**. Therefore, MIPergic LN activation could plausibly cause both fast-acting and long-lasting inhibition in the same and/or disparate downstream target. Moreover, MIPergic LN-derived GABA and MIP may simultaneously act on the same downstream target(s) to synergistically modulate their activity to have a greater effect than either modulator alone might achieve. Alternatively, MIPergic LNs might primarily use GABA throughout the course of ongoing network activity, and use MIP only under special circumstances (see example below). We attempted to parse the contribution of MIPergic LN-derived MIP from MIPergic LN-derived GABA by first determining what was the minimal strength of activation necessary to mobilize MIPergic LN dense core vesicles (DCVs). More specifically, we artificially activated MIPergic LNs by P2X2 misexpression^112^ and ATP injection, while simultaneously recording DCV changes via either ANF-GFP^48^ or a neuropeptide release sensor (NPPR-ANP-GCaMP6s^113^). However, we were unable to detect any change in either indicator even when we injected 100mM ATP (a concentration 10x-greater than what is necessary to activate other AL LNs^114^). As a result, it remains infeasible to simply artificially activate MIPergic LNs, while measuring a downstream neurons’ responses, and accurately attribute changes in the downstream neuron MIP released from MIPergic LNs. That said, because co-transmitters often increases the computational capacity a neuron and the plasticity of the networks in which they act^115–118^, how each specific MIPergic LN co-transmitter contributes to the overall role of these interneurons in AL processing is an important remaining question.

### Intrinsic and behavioral contributions of MIPergic modulation within the AL

Generally, multiple glomeruli are activated by any given odorant^25,53–55,119^. However, “optogenetic odors” can be used to selectively activate individual glomeruli in a manner similar to their odor-evoked responses to evaluate the behavioral contribution of individual glomeruli^120^. Such experiments reveal that DM1 and DM2 co-activation do not summate, and co-stimulation of both glomeruli produces a behavioral response that resembles DM1-only activation^120^. Based on this, the existence of an antagonistic relationship between DM1 and DM2 was proposed, wherein co-stimulation reduces the efficacy of either or both glomeruli^120^. We find MIP indirectly increases DM1 and directly decreases DM2 OSN responses **(Fig. 8)**. Therefore, MIP-SPR signaling in DM1 and DM2 may act as a homeostat such that coactivation of each glomerulus never produces a behavioral response greater than the DM1-only activation response. This “buffer” would be advantageous for preventing saturation at the downstream neurons that receives convergent input from these glomeruli^28,70,121,122^.

MIP-SPR signaling has been implicated in several behavioral state switches^23,24^. Notably, abolishing MIP release by inactivating all MIPergic neurons, or using a MIP-genetic null mutation, increases the animal’s drive for food-derived odors^23^. Moreover, DM2 OSN firing rate increases when all MIPergic neurons are inactivated^23^. In contrast, increasing the activity of all MIPergic neurons decreases attraction toward food-odors, to the extent of eliciting odor-induced aversion^23^. Together, these behavioral results suggest MIP-SPR signaling can affect the sensitivity to food-associated odors and drive to search for food. In accordance with these observations, we found that individual MIPergic LNs significantly co-innervate several food-odor associated glomeruli **(Fig. 2)** and neurons from several of these glomeruli express SPR **(Fig. 7)**. Most strikingly, we find that MIP directly acts on DM2 OSNs to decrease their odor-evoked responses **(Fig. 8)**. Furthermore, we show that the MIP-induced decrease in DM2 responses occurs in a stimulus-concentration independent manner **(Figs. 1 and 8)**. Altogether, these results point to a probable role for MIPergic LN-derived MIP signaling to adjust olfactory processing, likely while other MIPergic neurons adjust other sensory/motor elements, in accordance with satiety homeostasis drives. However, this role is likely only one of many that the MIPergic LNs play in AL processing as they also release GABA, and form reciprocal connections with neurons outside of the SPR-expressing neurons **(Fig. 3 and Fig. 4)**.

### Nuanced and non-intuitive emerging principles of peptidergic modulation

Peptidergic modulation can be as simple as a single neuropeptide modulating motor output in the stick insect locomotor system^123^, or as complex as the 37 neuropeptide families acting within the cortex^124^. Our data highlight how even a seemingly simple case, a single neuropeptide acting through a single receptor, can have complex consequences on network processing by acting non-uniformly within different components of the overall network. As neuropeptide functions are often deeply conserved, and as the actions of neuropeptides begin to come into focus, similar instances of complex and non-uniform peptidergic modulation will likely appear across disparate taxa and modalities.

## Methods

### Fly husbandry, genotypes, and subject details

A complete table of each animal’s genotype used for each experiment are included in **Supplementary Table 1**. Information on parental stock origins and relevant identifiers are provided in **Table 1**. Unless otherwise noted, flies were reared on standard cornmeal and molasses media at 24°C and under a 12:12 light:dark cycle. Equal numbers of male and female animals were used when possible, excluding live-imaging experiments which used only females. For mating status comparisons: 1) “virgin females” denotes females that were meconium-positive upon collection, 2) non-virgin females were housed with males until processing for immunohistochemistry, and 3) flies were age-matched and kept on the similar media until processed for immunohistochemistry.

**Table 1.**
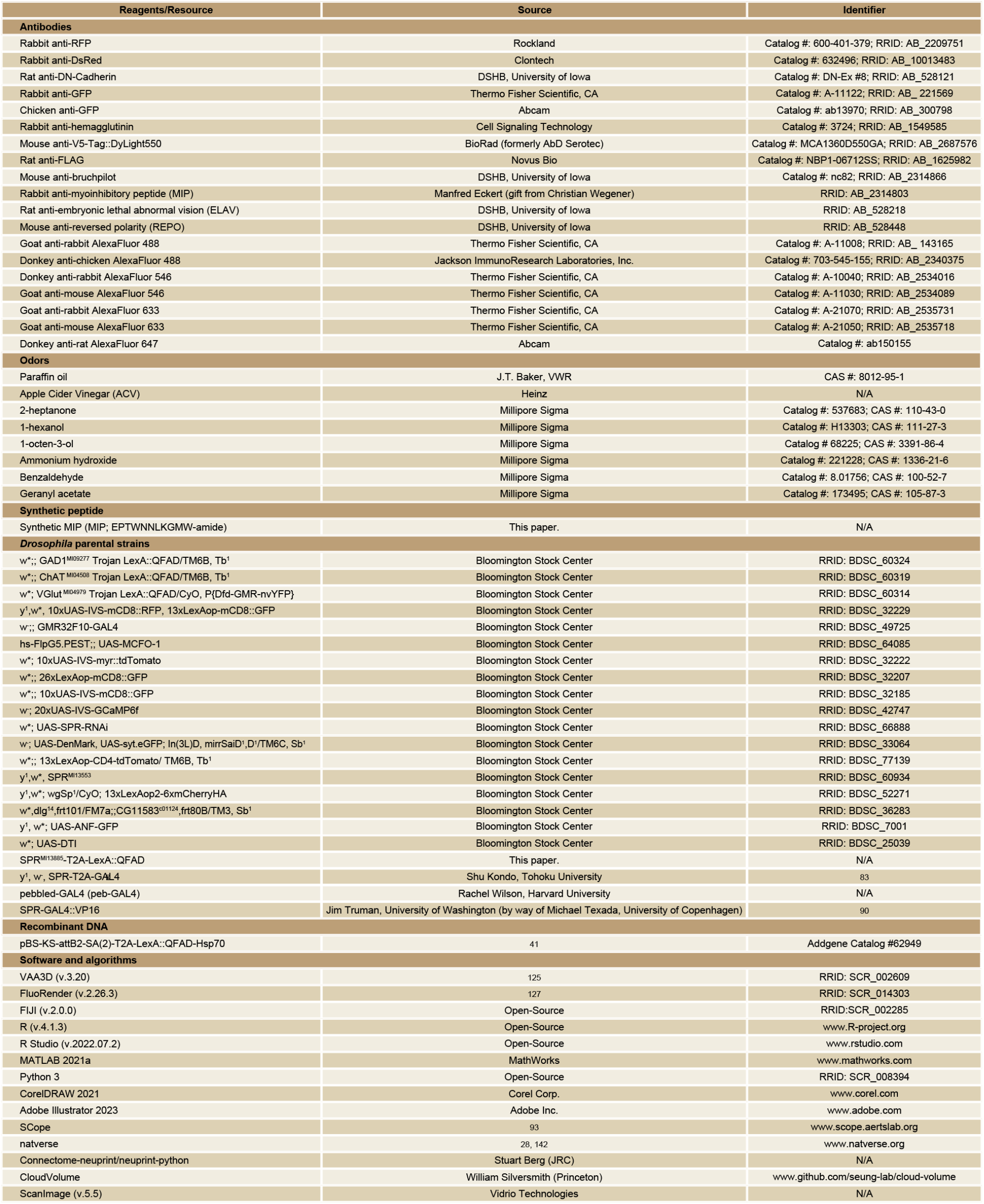
Sources and identifiers for all key reagents and resources used in this present study.

### Immunohistochemistry and imaging

All immunohistochemistry was performed generally as previously described^125^. Briefly, samples were dissected, fixed in 4% paraformaldehyde, then washed with phosphate buffered saline with 0.5% Triton-X 100 (PBST) several times before taking samples through an ascending-descending ethanol was series, then blocking in 4% IgG-free BSA (Jackson Immunoresearch; Cat#001-000-162). Samples were then incubated in primary antibody **(Table 1)** diluted in blocking solution and 5mM sodium azide. Following primary antibody incubation samples were washed with PBST, blocked, and incubated in secondary antibody diluted in blocking solution and 5mM sodium azide. Finally, samples were washed, cleared using an ascending glycerol series (40%, 60%, 80%), and mounted on well slides in Vectashield (Vector Laboratories, Burlingame, CA; Cat#H-1200). Images were collected and analyzed as previously described^125^ with VAA3D^126^ and FluoRender^127^, apart from those captured with a 40x/1.25 Silicone UPlanSApo Olympus objective.

### Single LN clone induction and glomerular innervation analyses

Single LN clones were induced through the MultiColor Flip Out (MCFO) method^49^. Flies carrying the MCFO cassettes, Flp-recombinase, and GAL4 driver were raised under normal conditions (see above) until heat shock. Adult flies were heat-shocked in a 37°C water bath for 12-25 minutes and returned to normal conditions for ~2-3 days before processing for immunohistochemistry. We chose to analyze the innervation patterns of 50 individual MIPergic LNs based on a statistical probability theorem termed, “the coupon collector problem”^128^. For our purposes, this meant we needed to sample 43 individual LNs to ensure we sampled each of the ~13 LNs highlighted by R32F10-GAL4 **(Fig. 1b-1d)**. We chose to analyze more than the minimal number as determined by this theorem as an additional preemptive measure to ensure the ~8 MIPergic AL LNs were sampled. Apart from VA1v, glomeruli were defined according to previously published AL maps^129,130^. Glomerulus names were later updated according to recent naming conventions^28^. Neuropil were labeled using anti-DN-cadherin or anti-Bruchpilot **(Table 1)**. Hierarchical clustering and principal components analysis (PCA) of glomerular innervation data were performed as previously described^32^. PCA was performed without any arbitrary threshold of significance. Glomeruli and individual MIPergic LN clones were hierarchically clustered using Ward’s method (“ward.D2”) and Euclidean distance using the “Heatmap” function in the *ComplexHeatmap* package^131^. Pairwise Pearson’s correlation coefficient for all possible binary combinations of glomeruli were determined from our MIPergic LN glomerular innervation clonal analysis data using the “cor” function in the base-R *stats* package. These Pearson’s correlation coefficients were subsequently assessed for statistical significance by using the “rcorr” function in the *Hmisc* package, which computes a matrix of Pearson’s r rank correlation coefficients for all possible pairwise combinations within a data matrix. The p-values in this instance are the probability that we would have found a given result if the correlation coefficient was zero (the null hypothesis). This is an indication of whether the aforementioned co-innervation of a given pair of glomeruli is significant or not. In other words, if “glomerulus A” and “B” are likely co-innervated by a given MIPergic LN (i.e., positive correlation of MIPergic LN between “glomerulus A” and “B”), is this likelihood statistically probable?” Further details regarding how significant correlations are computed using this approach are provided in the package’s documentation (https://github.com/harrelfe/Hmisc/blob/master/R/rcorr.s).

The *corrplot* package was used to create the hierarchically clustered (using Ward’s method) representation of these pairwise correlation coefficients depicted in **Fig. 1**. In every case used, glomerular “odor scene” information is derived from previous assignments^132^.

To determine if MIPergic LNs preferentially innervate glomeruli based on valence, glomeruli were assigned “attractive” or “aversive” based on similar assignments previously described^28,132^. These glomerular valences aggregate findings from previous reports^25,133–137^, as well as behavioral valence of the odors^138^ that glomerulus’ OSNs respond to according to DoOR 2.0^139^. Glomeruli whose valence is statedependent (e.g., the V glomerulus)^140^ and DC4 were not included in this analysis. Similar methods were used to determine if MIPergic LNs preferentially innervate glomeruli based on the functional group of a given OR’s cognate odorant, with the exception of the V and VM6 glomeruli.

### MIPergic LN ablation experiments

To determine whether MIPergic LNs are the sole source of MIP immunoreactivity within the AL, we used R32F10-GAL4 to drive the expression of a temperature-sensitive variant of Diphtheria toxin (UAS-DTI)^141^ in all MIPergic LNs. Animals carrying both transgenes were raised at a permissive temperature of 18°C, until ~2 days post-eclosion when they were shifted to the non-permissive temperature of ~25-28°C for ~3 days. After ~3 days at 25°C or 28°C animals were processed for MIP-immunolabeling as described above.

### MIPergic LN anatomical marker density analyses

Analysis of syt.eGFP, DenMark, anti-MIP immunoreactive puncta signal, and LN innervation density in antennal lobe glomeruli (via mCD8::GFP signal) was performed as previously described^75^. Images of all antennal lobes within a given brain were collected with similar confocal scan settings (laser power, detector offset, etc.) and later imported into FIJI for quantification. Using the Segmentation Editor plugin and a previously described script (graciously provided by Rachel Wilson, Harvard)^75^, ROIs were manually traced every 2-3 slices around the neuropil boundaries of each glomerulus using the anti-DN-Cadherin or anti-Bruchpilot channel, and then interpolated through the stack to obtain boundaries in adjacent slices. To ensure each brain contributed equally when pooling data across brains, signal density values for all glomeruli were normalized to the maximum density value within the given indicator being analyzed (e.g., all density values for syt.eGFP were normalized to the maximum syt.eGFP value). Signal density values were similarly normalized within-indicator, but also within-sex, for assessing sexual dimorphism in MIPergic LN syt.eGFP or DenMark puncta signal **(Supplementary Fig. 7)**. The “ggscatter” function in the *ggpubr* package was used to determine Pearson’s correlation coefficients and p-values when assessing correlations between effector/anti-MIP and MIPergic LN mCD8::GFP voxel density across all glomeruli. Adjusted R^2^ values were calculated using the base-R *stats* package and correspond to how well each data being assessed for the given correlation analysis fit a linear model.

### Putative MIPergic LN connectomic analyses - identifying putative MIPergic LNs

All connectome analyses leveraged the publicly available Janelia FlyEM *Drosophila* hemibrain electron microscopy volume (v.1.2.1; https://neuprint.janelia.org/)^60,61^, and recently described analysis suites^28,142^. We used several stringent criteria for determining which neurons are most likely MIPergic LNs, the first of which was the candidate neurons must be AL LNs. We then selected those candidate AL LNs that were previously determined to most likely belong to the patchy AL LN subtype^28^. Candidates were then filtered for those that receive input from the contralaterally projecting, serotonin-immunoreactive deutocerebral (CSD) neurons as all MIPergic LNs express the 5-HT1A serotonin receptor^125^, and form connections with the serotonergic CSD neurons^143^. We then used natverse^142^ to transform the interconnectivity of each candidate neuron into the FlyCircuit whole brain (FCWB) template brain three-dimensional space^144,145^, so we could generate a morphological similarity score between our query neuron and neurons FlyLight project’s GMR-GAL4 repository^47^ by using the built-in NBLAST package *(nat.nblast)^145^*. We selected for only those candidates that achieved a GMR32F10-GAL4 NBLAST score of >0.80, which is greater than the >0.60 score necessary to consider the query neuron and GMR GAL4 neurons “identical twins”^145^. Lastly, any remaining candidate MIPergic LNs were filtered for those neurons that are considered “Traced”, the hemibrain’s highest level of tracing completeness and confidence. Only neurons that met all of these criteria (~5% of all AL LNs) were considered for further analysis.

### Putative MIPergic LN connectomic analyses - putMIP LN meshes, segregation indices, and flow centrality

Most methods for analyzing putMIP LN morphology and connectivity have been described recently^28^. Putative MIPergic LN skeleton meshes **(Fig. 3g)** were fetched from the hemibrain data repository by accessing the neuPrint Python API using the *neuprint-python* (https://github.com/connectome-neuprint/neuprint-python) and *Cloud-Volume* (https://github.com/seunglab/cloud-volume) packages. The *hemibrainr* package (https://github.com/flyconnectome/hemibrainr) was used to fetch each putMIP LN’s metadata and calculate each neuron’s dendrite-axon segregation index and flow centrality^68^ using the recommended arguments.

### Putative MIPergic LN connectomic analyses - intraglomerular input:output ratio analysis

Glomerular meshes based on PN dendrites were used for all subsequent analyses (input:output ratio by glomerulus, connectivity demographics, etc.)^28^. To establish a input:output ratio for each glomerulus a given putMIP LN innervates, we extracted the number of input and output connections each putMIP LN has within each glomerulus by subsetting the connectors read in from the neuPrint database via the *neurprintr* “neuprint_read_neurons” function. These connectors were then filtered for their presence inside each glomerulus’ mesh XYZ coordinate space, segregated based on connection type (e.g., output), then finally summed. By analyzing the data in this manner, as opposed to simply considering the number of putMIP LN axons/dendrites within a given glomerulus, this analysis more likely closely captures putMIP LN input-vs.- output across the AL as *Drosophila* synapses are generally polyadic (reviewed in^62^). To establish a given putMIP LN’s input:output ratio across all glomeruli, we used the following formula: *(# of input connections - # of output connections)/(# of input connections* + *# of output connections).* Therefore, values from −1 to 0 indicate the given putMIP LN sends more output within the given glomerulus. Conversely, values from 0 to 1 indicates the given putMIP LN receives more input within the glomerulus.

### Putative MIPergic LN connectomic analyses - general upstream and downstream demographics analyses

To identify and compare the demographics of each putMIP LN’s upstream and downstream partners, putMIP LN connectivity data were first extracted using the *hemibrainr* “simple connectivity” function. The demographic of each presynaptic and postsynaptic partner was generally assigned according to the neuron’s accompanying “name” or “type” as listed on neuPrint, or by previously established cell type assignments^28^. In cases where a neuron’s “name” or “type” was unannotated (“NA”), the neuron would be categorized as “Unknown”. We used the following formula to determine the percentage of overall input a given putMIP LN receives from a given neuron category: *[(sum of connections from a given neuron category to the given putMIP LN)/(summed amount of input that given putMIP LN receives from all categories)] x 100%*. Similar methods were applied for determining the percentage of overall output a given neuron category receives from a given putMIP LN.

### Putative MIPergic LN connectomic analyses - putMIP LN input polarity analysis

To determine the amount of excitatory, inhibitory, and modulatory input a given putMIP LN receives within each glomerulus, we first categorized each presynaptic neuron as either excitatory, inhibitory, or modulatory based on the presynaptic neuron’s neuPrint “name”/”type”, previous immunohistochemistry results^31,32,35,146–149^, and/or the category assigned in previous reports^28^. However, we acknowledge several caveats to this analysis, such as: (*1*) this analysis does not account for co-transmission; (2) several glomeruli are truncated within the hemibrain AL^28^; (3) although we consider all LNs as inhibitory as most are either GABAergic or glutamatergic (combined, these represent ~170/200 AL LNs)^31^,^32,35,114,146,148^, there are ~4 tyrosine hydroxylase-immunoreactive (dopaminergic) and ~8-15 cholinergic and/or electrically coupled LNs in the AL^32,33,38,39^; (4) although GABA can also act as an intrinsic modulator in the AL (reviewed by Lizbinski & Dacks^150^), we only count GABAergic LNs as part of the “inhibitory input” category here; and, (5) we consider all ventral LNs analyzed here as being glutamatergic, but there are ~4 dopaminergic (tyrosine hydroxylase-immunoreactive) ventral LNs^32^. Once each presynaptic neuron’s chemical identity (excitatory, inhibitory, modulatory, or unknown) was determined, we used several approaches to assign these synapses to particular glomeruli. In the case of uniglomerular PNs (uPNs) and OSNs, we leveraged the single glomerulus innervation of these presynaptic neuron types to assign their synapse onto a given putMIP LN synapse to the presynaptic neuron’s home glomerulus. That is to say, OSN-to-putMIP LN and uPN-to-putMIP LN synapses were assigned to a glomerulus by: (*i*) using the home glomerulus assigned to a given presynaptic in the neuron’s neuPrint “name”/”type”, or (*ii*) by the home glomerulus assigned to the neuron in previous reports^28^. For instance, if the presynaptic neuron was a cholinergic PN whose home glomerulus is DA2, and this DA2 PN synapses on a given putMIP LN five times, then those five synapses went to the overall excitatory input the given putMIP LN receives within DA2. Neurons were only excluded from this analysis if the presynaptic neuron’s home glomerulus was not previously identified^28^. Once the polarity of the input type was established, we used the same methods as above for determining whether the XYZ coordinates of each putMIP LN’s synapse(s) with a given presynaptic partner were located in a given glomerulus. Synapse counts for each putMIP LN partner within the given glomerulus were then summed by type (excitatory, inhibitory, modulatory, or unknown), and the resulting total was divided by the total number of synapses the given putMIP LN makes within that glomerulus to establish percent excitatory, inhibitory input, or modulatory input.

### SPR^MI13885^-T2A-LexA::QFAD generation

The SPR^MI13885^-T2A-LexA::QFAD fly line was established using previously described injections methods^41^. We also note that we also attempted to create an SPR-T2A-GAL4 using the pC-(lox2-attB2-SA-T2A-Gal4-Hsp70)3 construct (Addgene #62957), but no founders emerged (potentially owing to lethality when these construct elements are inserted in the SPR locus). Briefly, pBS-KS-attB2-SA(2)-T2A-LexA::QFAD-Hsp70 and ΦC31 helper plasmid DNA were co-injected into y^1^, w*, MiMICSPR^MI13885^. pBS-KS-attB2-SA(2)-T2A-LexA::QFAD-Hsp70 (Addgene plasmid #62949) and pC-(lox2-attB2-SA-T2A-Gal4-Hsp70)3 (Addgene #62957) were gifts from Benjamin H. White (NIH). SPR^MI13885^-T2A-LexA::QFAD transformants were isolated as depicted in **Supplementary Fig. 7**.

### Single-cell RNA-sequencing (scRNA-seq) analysis of SPR expression

Single-cell transcriptomic data were accessed and downloaded from the SCope web interface (https://scope.aertslab.org) on 03/04/2022. Projection neuron clusters were re-identified as in each dataset’s original report^91–93^. Transcript reads were exported log-transformed (log(1 + x)) and reads were counts-per-million (CPM) normalized. Projection neuron subpopulations were then identified within each scRNA-seq dataset using previously established marker genes^91,151,152^.

### *in vivo* calcium imaging - animal preparation

All calcium imaging experiments were performed on female flies ~1-5 days post-eclosion, and at room temperature. All physiology occurred within the animal’s ZT0 and ZT8. Animals of the proper genotype were collected and briefly anesthetized on ice. Once anesthetized, an animal was affixed to a custom-built holder with UV curable glue (BONDIC, M/N: SK8024). Our custom-built holder consists of a sheet of aluminum foil with a ~1×1mm square (the imaging window) affixed to a 3D-printed design derived from similar designs described previously^153^. Once mounted, a small window exposing the dorsal side of the brain was created, and covered with twice-filtered recording saline (in mM: 2 CaCl_2_, 5 KCl, 5 HEPES, 8.2 MgCl_2_, 108 NaCl, 4 NaHCO_3_, 1 NaH_2_PO_4_, 10 sucrose, and 5 trehalose; adjusted pH: ~7.4)^29^. After establishing the imaging window, the air sacs, fat bodies, and trachea covering the dorsal side of the brain - as well as Muscle 16 - were removed with fine forceps. With the exception of minimal epochs during the synthetic MIP bath application experiments (see below), the brain was continuously perfused with oxygenated (95%O_2_/5%CO_2_) recording saline using a Cole-Parmer Masterflex C/L (M/N: 77120-62) at a rate of ~2mL/min.

### *in vivo* calcium imaging - image acquisition

For one-photon imaging data (the majority of *in vivo* physiology data), data were acquired using a Prior Scientific Open Stand (M/N: H175) microscope mounted on Prior Scientific motorized translational stage (M/N: HZPKT1), and equipped with an Olympus 10x/0.30 UPlanFL N objective and an Olympus 60x/1.00 LUMPlanFL N water-immersion objective. A 470nm CoolLED pE-100 (CoolLED Ltd., Hampshire, UK) was used as the light source. Each trial was captured with a Hamamatsu ORCA-4.0LT camera (Hamamatsu Phototonics, Hamamatsu, Japan), and consists of 40 1,024×1,024 frames acquired at a frame rate of ~9 Hz.

A portion of the R32F10-GAL4 odor panel experiments were also acquired using a custom-built two-photon system (Scientifica) equipped with a Mai Tai HP Ti:Sapphire laser (Spectra-Physics) and operated using ScanImage acquisition software (v.5.5; Vidrio Technologies). Emitted fluorescence was captured by a gallium arsenide phosphide (GaAsP) photomultiplier-tube detectors. Each trial consisted of 80 512×512 frames acquired at a frame rate of ~3.4 Hz. After data acquisition, a high-resolution z-stack (1,024×1,024) was acquired at ~0.21 Hz to enable post-hoc glomerulus identification as previously described^29,40,154–156^ (also, see below).

### *in vivo* calcium imaging - odor preparation and delivery

All odor concentrations are reported as v/v dilutions in paraffin oil (J.T. Baker, VWR #JTS894), or autoclaved and twice-filtered distilled water (for diluting acids). For example, 10^-2^ dilution indicates that one volume of an odor is diluted with 100 volumes of paraffin oil. For one-photon imaging data (the majority of *in vivo* physiology data), dilutions were prepared in 2mL odor vials (SUPELCO; P/N: 6020) that contained a final volume of 1mL of diluted odor in paraffin oil every other day, or after two experiments (whichever came first). Odors were generally presented as previously described^75,77,122^. Briefly, a carrier stream of carbon-filtered, dehumidified, air was presented at 2.2 L/min to the fly continuously through an 8mm Teflon tube placed ~1cm away from the fly. A three-way solenoid (The Lee Company, P/N: LHDA1231315H) diverted a small portion of the airstream (0.2 L/min) through the headspace of an odor vial for 200ms after triggering an external voltage command (TTL pulse) at frame 20 of the trial. Considering the above, the odor is diluted further (by 10-fold) prior to delivery to the animal. The odor stream joined the carrier stream 11cm from the end of the tube, and the tube opening measured ~4mm. Odor delivery during two-photon imaging was similar, but differed slightly in that: (1) odor cartridges (see below) instead of a 2mL odor vial; (2) the continuous airstream was presented via a custom-built glass tube; and, (3) the TTL pulse occurred at frame 30 of the trial.

Methods for assessing preparation health and performing multiple odor trials conform to previous work^75,122^. At the start of each experiment, the animal was presented a test odor (10^-3^ 2-heptanone) to assess the preparation’s health. Only the data collected from animals whose responses to this test odor were robust and did not dramatically change from baseline over the course of the experiment were used for further analysis. The only exceptions to this were those data collected in synthetic MIP bath application experiments (see below), since bath application of any modulator would likely result in network property changes that would consequently change olfactory responses. Therefore, the test odor was only initially presented to those animals used for synthetic peptide application experiments, so their initial olfactory response health could be assessed. Each experiment consisted of multiple odor trials (3 for OSNs; 4 for LNs) within a preparation which were then averaged to attain a within-animal response. These within-animal averages were subsequently averaged across many animals for subsequent statistical analysis, and “n” is reported as the number of animals. Each odor trial consisted of five 200ms pulses of odor with a 1ms interpulse interval. The same odor was never presented twice within 2min to prevent depletion of the odor vial’s headspace. If multiple odors were to be tested, then they were presented randomly. If multiple concentrations of a given odor were to be tested, then the lower concentration was presented before the higher concentration. Air entered and exited each odor vial through a PEEK one-way check valve (The Lee Company, P/N: TKLA3201112H) connected to the vial by Teflon tubing. The odor delivery tube was flushed with clean air for 2min when changing between odors/concentrations. As an additional preemptive measure, all odor delivery system components were hooked up to the house vacuum line overnight. The olfactometer used in two-photon data collection consisted of odor cartridges (a syringe housing a piece of filter paper that was doused in 10μl of diluted odor) hooked into a custom glass carrier stream delivery tube as previously described^157^.

### *in vivo* calcium imaging - data analysis

All calcium imaging data were analyzed using a custom-made MATLAB script graciously provided by Marco Gallio (Northwestern University) and has been described previously^51,158,159^. With the exception of any preparations that violated the aforementioned criteria (e.g., movement, diminishing prep health, etc.), no data points or outliers were excluded from our analyses. Generally, the number of flies to be used for experiments are not a limiting factor, therefore no statistical power analyses were used to pre-determine sample sizes. Regardless, our sample sizes are similar to those in previous reports that perform similar experiments^30,51,160–165^. Before analyzing the data, a Gaussian low-pass filter (sigma=1), bleach correction (exponential fit), and image stabilizer algorithms were applied to the given trial’s raw ΔF/F signal. Similar preprocessing for two-photon microscopy data was similar, with the exception of a higher sigma during Gaussian low-pass filtering (sigma=2). A trial’s average fluorescence image was used as a guide to draw consistently sized circular regions-of-interest (ROI) within a given glomerulus. Calcium transients (ΔF/F) within the ROI were measured as changes in fluorescence (ΔF) normalized to baseline fluorescence (F, fluorescence intensity averaged across 2sec just prior to odor onset). Within-animal responses were established by averaging across several odor trials in the given preparation (3 for OSNs; 4 for LNs). These within-animal responses were then pooled for each stimulus identity and concentration across animals. These pooled averages were used for all subsequent statistical analyses and the “n” is reported as the number of animals. Glomeruli were manually identified post-hoc by comparing acquired images to well-defined three-dimensional maps of the AL^166,167^. Only the glomeruli that were reasonably identifiable were considered for analysis.

### Myoinhibitory peptide (MIP) application experiments

MIP (MIP; EPTWNNLKGMW-amide) was custom made by GenScript (Piscataway, NJ, USA) at the highest purity available (>75%). The sequence we chose to use for MIP is identical to the sequence previous investigations have used when discerning the role of MIP in the Drosophila circadian system^161^. In pilot experiments, we tested another sequence of MIP (RQAQGWNKFRGAW-amide) that was previously detected at the highest abundance by direct profiling of single ALs using mass spectrometry^26,168^. Experimental results produced using peptide of either sequence were not qualitatively different, but all results reported here use the MIP previously used in circadian studies^161^. To test how MIP application adjusts odor-evoked responses, a 1,000μM working solution was made by diluting a small portion of the lyophilized peptide in nuclease-free water (ThermoScientific, #R0581). After testing the initial odor-evoked responses of the neurons being tested for a given experiment, the perfusion system was momentarily switched off so a small portion of our MIP working solution could be pressure injected into the AL to a final concentration of 10μM. This final concentration was chosen for several reasons, which include: (1) we wished to remain consistent with other studies of peptidergic modulation in the *Drosophila* AL^29,30,160^; (2) we wished to be consistent with studies on the effects of MIP in other circuits^161^; and, (3) previous reports have already determined that our chosen effective concentration (10μM) is the optimal concentration for testing the effect of MIP on *Drosophila* neurons^161^. Ten minutes after MIP pressure injection, the animal’s odor-evoked responses were tested as before MIP injection, and then the perfusion system was switched back on. Ten minutes after turning the perfusion system back on, the animal’s odor-evoked responses were once again tested as they were initially. Re-testing the animal’s response to the test odor (10^-3^ 2-heptanone) at the end of these experiments could not be used as a reliable means for assessing prep health due to changes in circuit member responses induced by modulator bath application. Therefore, for these experiments no animal was tested for longer than the average time that animals were reliably healthy in the MIPergic LN odor panel experiments (~90min). Furthermore, we believe these preparations remain healthy throughout the entire experimental epoch as ACV responses increase or do not significantly diminish over the course of the experimental epoch in many glomeruli **(Fig. 1)**.

### Quantification and statistical analyses

#### General approach

Statistical analyses were performed using R (v.4.1.3) in R Studio (v.2022.07.2). Values to be analyzed were concatenated in Excel before importing into the relevant analysis software. Statistical results are reported within the main text and/or figure legends. All statistical tests were twotailed. All boxplots display the minimum, 25^th^-percentile, median, 75^th^-percentile, and maximum of the given data. Additional analysis details are provided for each set of experiments above. Where possible, values are given as mean ±SEM. Statistical significance is defined as: *p ≤ 0.05, **p ≤ 0.01, ***p ≤ 0.001.

#### Statistical analyses related to neuroanatomical experiments

The *ComplexHeatmap* package was used to hierarchically cluster glomeruli and individual MIPergic LN clones using Ward’s criteria and Euclidian distance. The *ClustVis* package (https://github.com/taunometsalu/ClustVis)^117^ was used to perform PCA on individual MIPergic LN innervation patterns. The “cor” function in the base-R *stats* package and the “rcorr” function in the *Hmisc* package were used to calculate statistically significant Pearson’s correlation coefficients for MIPergic LN pairwise glomerular innervation patterns. The *ggpubr* package’s “ggscatter” function was used to determine Pearson’s correlation coefficients and p-values when assessing correlations between: (1) effector/anti-MIP and MIPergic LN mCD8::GFP voxel density across all glomeruli, and (2) MIPergic LN glomerular innervation frequency as a function of each glomerulus’ volume. Adjusted R-squared values were calculated using the base-R *stats* package and correspond to how well each data being assessed for the given correlation analysis fit a linear model. The Shapiro-Wilk test (the *rstatix* package’s “shapiro_test” function) was used to evaluate any deviations from a normal distribution. Welch’s unpaired t-test was used to determine if MIPergic LNs preferentially innervate glomeruli based on inferred hedonic valence. A Kruskal-Wallis rank sum test followed by pairwise Bonferroni’s-corrected Dunn’s multiple comparisons test was used to determine if: (1) MIPergic LNs preferentially innervate based on the functional group found along the odorant that activates the given glomerulus’ odorant receptor; (2) SPR-GAL4::VP16 expression in antennae and maxillary palps significantly differs between males, mated females, and virgin females; (3) SPR-GAL4::VP16 expression in glutamatergic LNs between males, mated females, and virgin females; (4) SPR-T2A-GAL4 expression in maxillary palps significantly differs between males, mated females, and virgin females; and, (5) the number of MIPergic LNs differ between males, mated females, and virgin females. Welch’s one-way ANOVA with a Bonferroni multiple comparisons correction was used to assess statistically significant differences in SPR-T2A-GAL4 expression in antennae between males, mated females, and virgin females. A two-way ANOVA with a Greenhouse-Geisser sphericity correction followed by a Bonferroni’s multiple comparisons test was used to assess sexual dimorphism in MIPergic LN syt.eGFP or DenMark puncta density across glomeruli.

#### Statistical analyses related to physiology experiments

Background-subtracted changes in fluorescence over time (ΔF/F) analyses were carried out using custom MATLAB scripts previously described^51,158^, and are represented as individual traces overlaid by the mean with dilutant-only (e.g., paraffin oil-only) responses subtracted. Peak response **(Fig. 5)** refers to the maximal ΔF/F value within the time of odor onset to ~1 second post-odor onset averaged across all animals. Area under the ΔF/F curve (AUC) was modified from previous reports^169^, such that AUC was calculated using Simpson’s rule (“sintegral” function in *Bolstad2* package) as the integral of the ΔF/F traces from the beginning until 1 second after odor delivery with a baseline of 1 second before stimulus onset. To assess OSN odor-evoked response differences across MIP treatments, we first determined if normality could be assumed (as above). If normality could be assumed, then an omnibus repeated measures one-way ANOVA with a Greenhouse-Geisser sphericity correction was performed (RM one-way ANOVA) (“anova_test” function in *rstatix).* If significant differences were detected with the omnibus, then pairwise repeated measures t-tests (RM t-tests) with a Holm multiple comparisons correction were performed to identify which groups were statistically different. If normality could not be assumed, then a Friedman rank sum test followed by Holm-corrected paired two-sided Wilcoxon signed-rank test was performed. All effect sizes reported were calculated using either the “cohens_d” (for parametric data) or “friedman_effsize” (for non-parametric data) function from the *rstatix* package, which compute Cohen’s d or Kendall’s W, respectively.

## Data availability

Connectomic and scRNA-seq source data are available on neuPrint (https://neuprint.janelia.org/) and SCope (https://scope.aertslab.org/), respectively. Any additional information required to reanalyze the data reported here is available from the lead contact upon reasonable request.

## Code availability

With the exception of code that was graciously provided to us by others, all code that was used to analyze or plot data is available from the lead contact upon reasonable request.

## Materials availability

Further information and reasonable requests for reagents and resources should be directed to and will be fulfilled by either Tyler R. Sizemore or Andrew M. Dacks. All novel transgenics generated here will be deposited with the Bloomington Drosophila Stock Center post-publication.

## Acknowledgments

We are grateful to Kristyn Lizbinski, Eric Horstick, and John Carlson for the helpful comments given on earlier manuscript drafts. Flies were kindly shared or acquired from: Shu Kondo, David Krantz, Michael Texada, Jim Truman, Rachel Wilson, Quentin Gaudry, the Bloomington Drosophila Stock centers (NIH P40OD018537), and the Janelia Fly Light project for flies. Christian Wegener kindly provided the MIP antibody that was developed by Manfred Eckert. Marco Gallio and Rachel Wilson graciously provided analysis scripts. Karen Menuz gave helpful insight in the interpretation of RNA-sequencing results from prior work. Stephen Plaza, Alexander Bates, Marta Costa, Greg Jefferis, and Philipp Schlegel gave helpful advice and instruction regarding connectomic analyses, and graciously shared odor scene and valence information. Our connectomic analyses also benefited from discussions in the nat-user community (https://groups.google.com/g/natuser), and a workshop organized by the Virtual Fly Brain and Drosophila connectomics teams (virtualflybrain.org; WT 208379/Z/17/Z). Fengqiu Diao gave technical advice for creating the SPR^MI13885^-T2A-LexA::QFAD Trojan exon transgenic animals. Kristyn Lizbinski and James Jeanne graciously provided advice and equipment specifications that lead to the construction and use of the olfactometer system described here. Kristyn Lizbinski, James Jeanne, Marco Gallio, Mehmet Keles, and Gaby Maimon provided invaluable technical advice for performing *in vivo* physiology. Kevin Daly kindly loaned equipment to us for our purposes and gave invaluable technical advice regarding their use. We would also like to thank Jacob Ralston for assisting by collecting confocal scans of AL anti-MIP labeling from R32F10-GAL4 diphtheria toxin ablation experiments. This work was supported by a Grant-In-Aid of Research (G20141015669888) from Sigma Xi, The Scientific Research Society (T.R.S.), start-up funds from WVU (A.M.D.), a National Institutes of Health R03 DC013997 (A.M.D.), a National Institutes of Health R01 DC016293 (A.M.D.), and two AFOSR DURIP awards (FA9550-19-1-0179 and FA9550-20-1-0098) (A.M.D.).

## Author contributions

T.R.S. conceived and implemented experiments, analyzed the subsequent data, prepared figures, and wrote and edited the manuscript. J.J. acquired data that contributed to Fig. 5. A.M.D. conceived experiments, acquired funding, and helped with manuscript edits.

## Supplementary information

**Supplementary Fig. 1.**
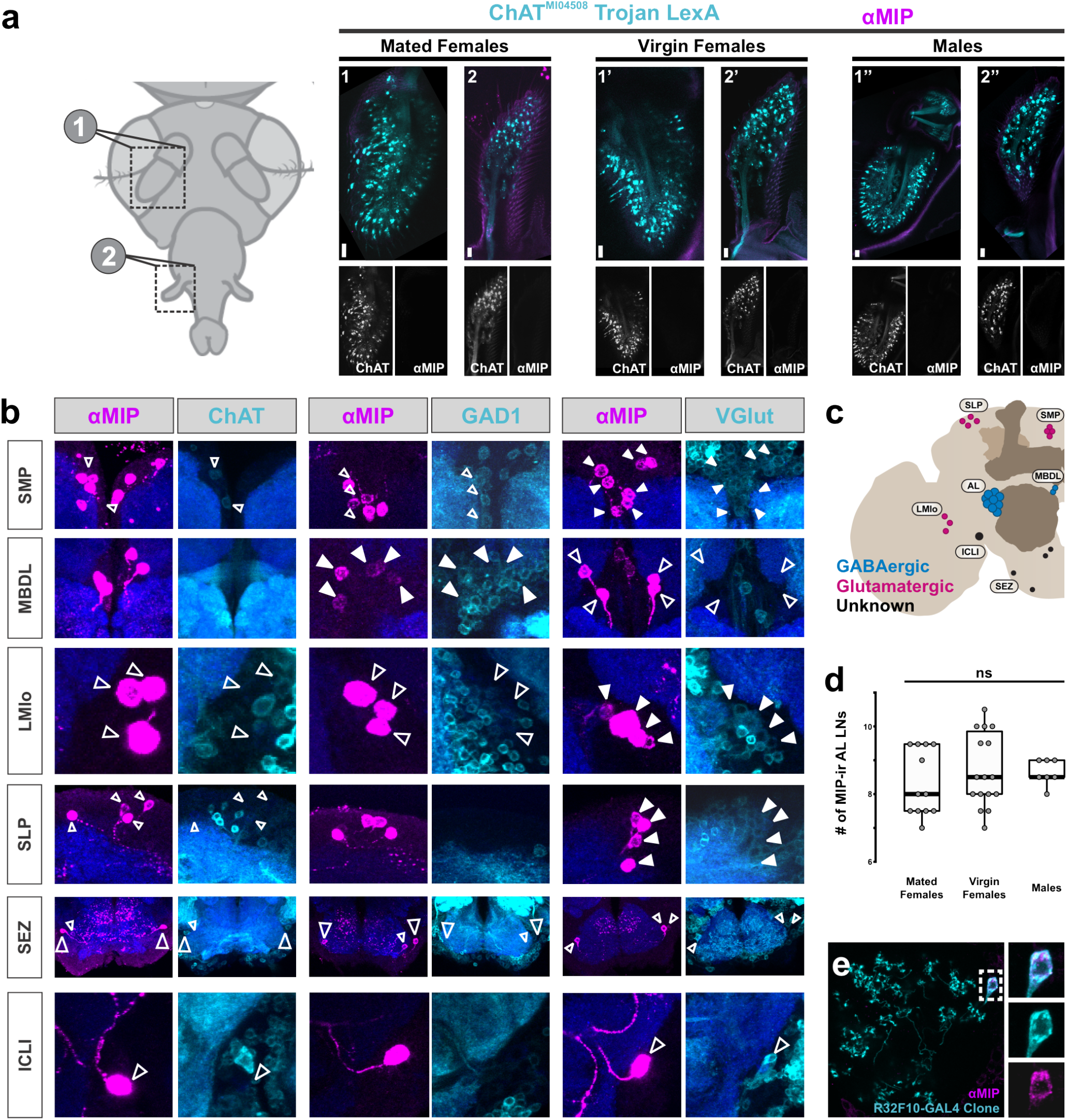
Myoinhibitory peptide (MIP) colabeling with transgenic markers for GABAergic, cholinergic, and glutamatergic neurons in the *Drosophila* central brain. **(a)** Regardless of sex or mating status, there are no MIP-immunoreactive (MIP-ir) OSNs in *Drosophila*. The left most diagram represents the imaging plane for all images to the right, wherein: **1-1”** = OSNs in the 3^rd^-antennal segment; **2-2”** = maxillary palp OSNs. High-contrast black-and-white images for each individual label (ChAT Trojan LexA-derived tdTomato or anti-MIP) are shown below each merged image (images in color). **(b)** MIPergic neurons in the antennal lobe (AL) **(see also Fig. 1)** and near the median bundle (MBDL) colabel with glutamic acid decarboxylase 1 (GAD1). MIPergic neurons in the superior medial and lateral protocerebrum (SMP and SLP, respectively) and near the lateral medial lobula (LMlo) colabel with vesicular glutamate transporter (VGlut). MIPergic neurons within the inferior contralateral interneuron cluster (ICLI)^170^ and SEZ do not colabel for ChAT, GAD1, or VGlut, and are most likely tyraminergic (Tyr) based on scRNA-seq data^92^. **(c)** Schematic summarizing data from **b**, wherein several populations of MIP-immunoreactive neurons are also glutamatergic (MIP^+^-VGlut^+^ neurons in the SMP, LMlo, and SLP; magenta), two populations are also GABAergic (MIP^+^-GAD1^+^ neurons in the MBDL and AL) **(see also Fig. 2)**, and no MIP-immunoreactive neurons are cholinergic (colabel with ChAT). Except for the ICLI interneurons, soma locations are labeled according to the closest neuropil, or fascicle, according to established nomenclature^171^. **(d)** The number of MIP-ir AL LNs does not differ based on sex or mating status (p = 0.548, Kruskal-Wallis rank sum test). Cell counts: n = 12 brains, 24 ALs (<u>mated female</u>); n = 16 brains, 31 ALs (<u>virgin female</u>); n = 7 brains, 14 ALs (<u>male</u>). **(e)** Representative image of R32F10-GAL4 clone colabeled for MIP shows MIPergic LNs are bonafide patchy AL LNs. In all cases: neuropil was delineated by anti-DN-cadherin staining; open arrowheads = no colocalization; closed arrowheads = colocalization; scale bars = 10μm.

**Supplementary Fig. 2.**
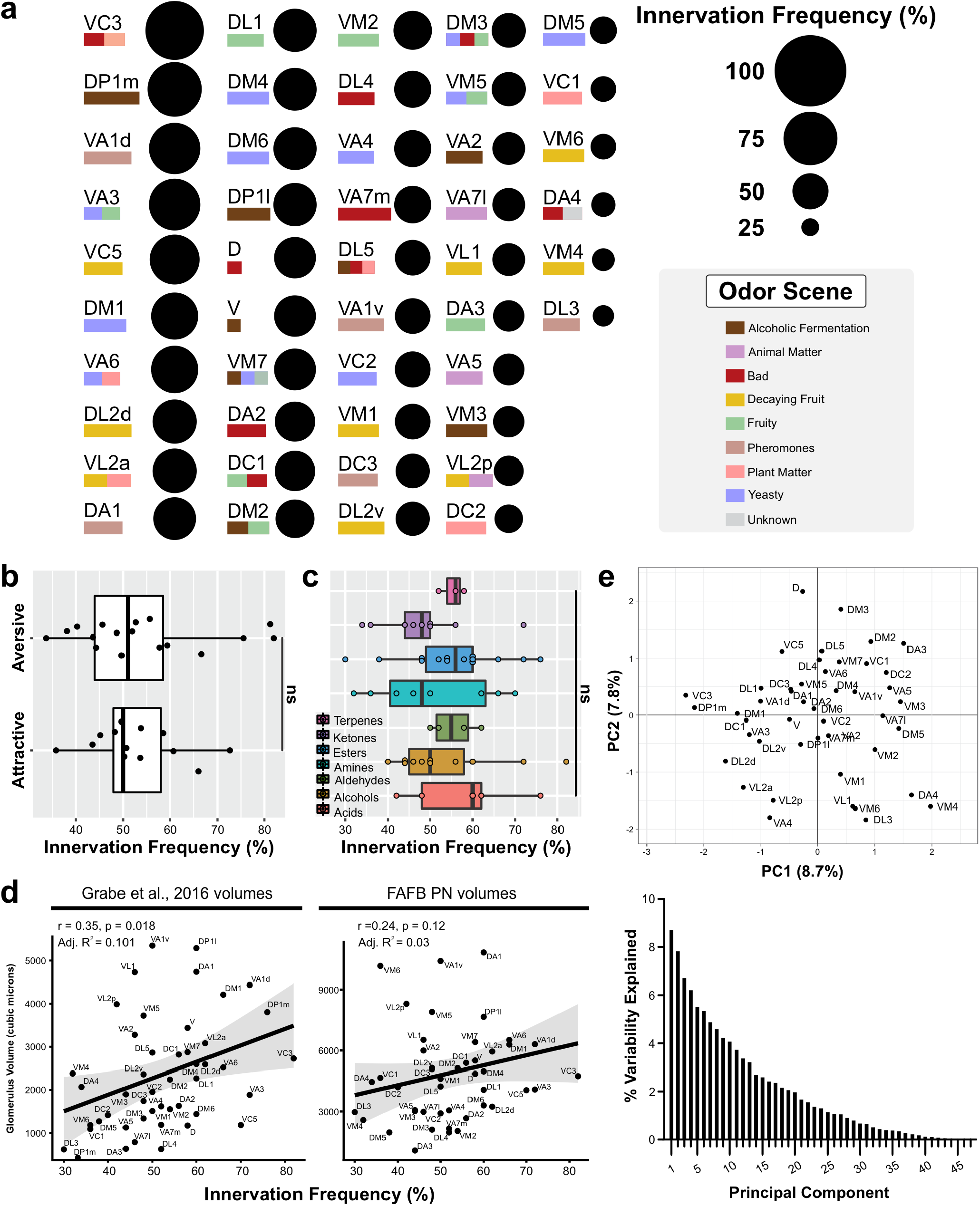
MIPergic LNs do not preferentially innervate olfactory glomeruli based on odor-evoked behavioral valence, the odor-tuning properties of a given glomeruli’s olfactory receptor neuron(s), or the size of the glomerulus. **(a)** Dot plot representation of the frequency we find a given glomerulus is innervated by a single MIPergic LN clone. Rectangles underneath each glomerulus’ name represents the “odor scene” of that glomerulus^28,132^. These are: alcoholic fermentation (brown); yeasty (blue); fruity (faded green); decaying fruit (yellow); plant matter (pink); animal matter (pale purple); pheromones (chartreuse); dangerous (red); and, unknown (grey). **(b)** MIPergic LNs do not preferentially innervate glomeruli whose activity has been linked to attractive or aversive behavioral responses (p = 0.991, n = 13 (“attractive”), 16 (“aversive”), unpaired t-test with Welch’s correction). **(c)** MIPergic LNs do not preferentially innervate glomeruli tuned to any particular odorant molecules (p = 0.59, Kruskal-Wallis rank sum test). Odorant molecule functional groups are color coded as follows: terpenes (magenta), ketones (purple), esters (blue), aromatics (aqua marine), amines (chartreuse), aldehydes (green), alcohols (brown), and acids (deep pink). **(d)** The frequency by which a MIPergic LN innervates a glomerulus is not correlated to the volume of the glomerulus (cubic microns). MIPergic LN innervation frequencies are significantly weakly correlated to glomerular volumes delineated by Grabe et al.^86^ (r = 0.35, p = 0.018), but variations in MIPergic innervation frequencies across glomeruli do not correlate (adjusted R^2^ = 0.101). Conversely, MIPergic LN innervation frequencies do not correlate with projection neuron-based glomerular volumes delineated from electron microscopy data^28,132^ (r = 0.24, p = 0.12). **(e)** Principal components analysis (PCA) of MIPergic LN innervation patterns, where each data point represents MIPergic LN innervation patterns for each glomerulus. Bar graph represents the percentage of the variance explained by each principal component.

**Supplementary Fig. 3.**
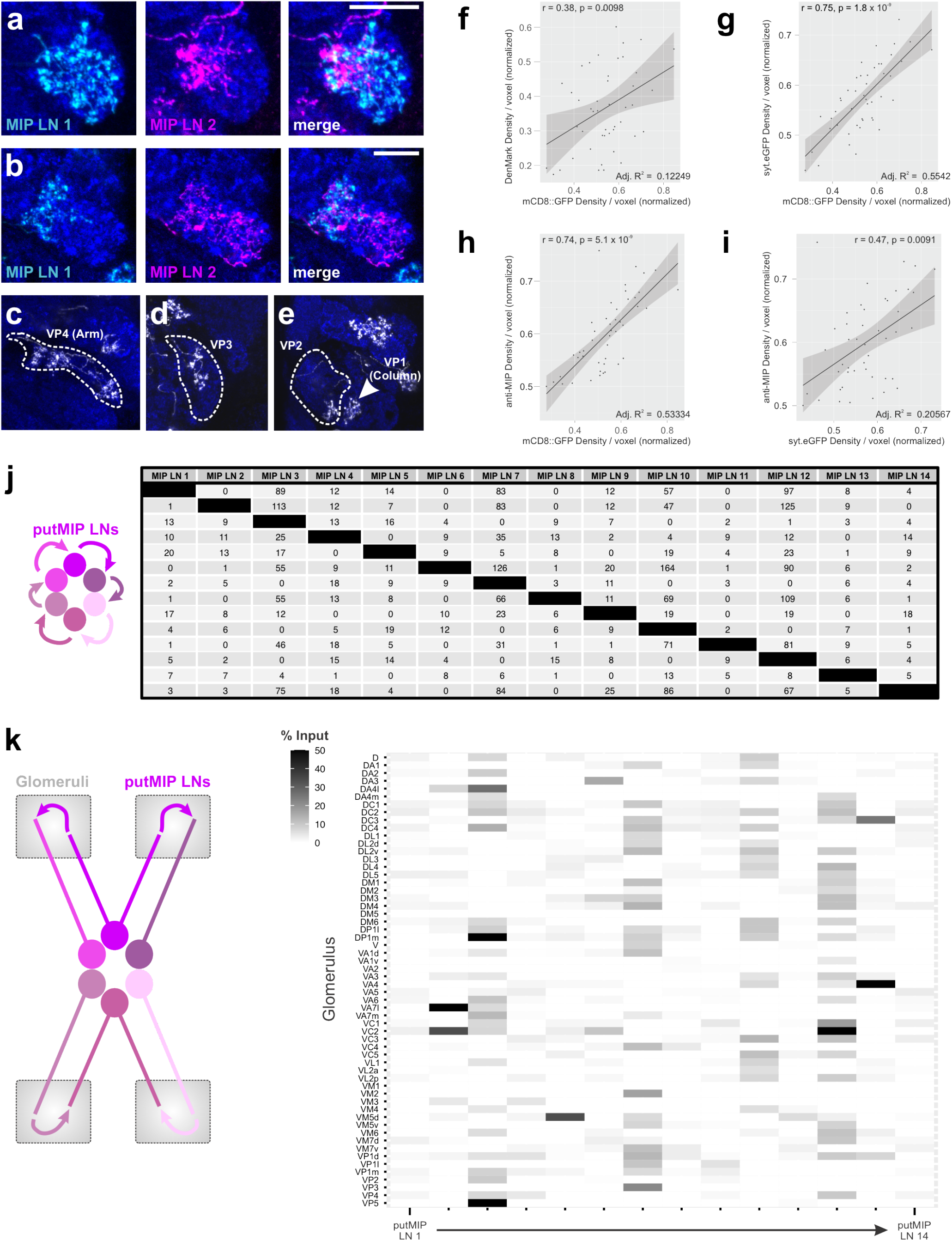
Sister MIPergic LN and individual MIPergic LN connectivity dynamics. **(a & b)** On average, ~12 glomeruli are co-innervated by sister MIPergic LN clones. In these examples, two distinct MIPergic LNs co-innervate DL2d and DP1l (respectively). For comparing sister MIPergic LN co-innervation patterns, n = 5 brains; 5 sister clones per brain. **(c-e)** Individual MIPergic LNs innervate thermo-/hygrosensory glomeruli. Branching from an individual MIPergic LN was observed invading the VP4 (formerly, “the arm”), VP3, VP2, and VP1 (formerly, “the column”). VP2-4 are designated by the hatched outline, while an arrowhead designates VP1. **(f-i)** DenMark, synaptotagmin-eGFP (syt.eGFP), and anti-myoinhibitory peptide immunoreactive puncta (anti-MIP) voxel density generally scale with MIPergic LN total cable voxel density within glomeruli. DenMark variations across glomeruli are significantly weakly correlated to the voxel density of total MIPergic LN cable within each glomerulus (r = 0.38, p = 0.0098). However, variations in DenMark voxel density across glomeruli do not correlate (adjusted R^2^ = 0.12249). Synaptotagmin-eGFP and anti-MIP voxel density are significantly correlated with the voxel density of total MIPergic LN neurite volume (<u>syt.eGFP</u>: r = 0.75, p = 1.8×10^-9^; <u>anti-MIP</u>: r = 0.74, p = 5.1×10^-9^). The density of syt.eGFP and anti-MIP immunoreactive punctate are significantly correlated (p = 0.0091), but variations in either indicator across glomeruli are not (adjusted R^2^ = 0.20567). In all cases, each data point represents the normalized mean indicator density within a given glomerulus and each line represents the linear regression model. **(j)** All putMIP LNs are synaptically connected to each other. Table of the number of synapses from one putMIP LN to all other putMIP LNs. **(k)** The amount of putative MIPergic LN reciprocal connectivity assessed within each glomerulus. Heatmap of the amount of input a given putMIP LN (x-axis) receives from all other putMIP LNs within every AL glomerulus as a function of the total amount of input that putMIP LN receives within a glomerulus. In all cases: neuropil was delineated by anti-DN-Cadherin staining; scale bars = 10μm.

**Supplementary Fig. 4.**
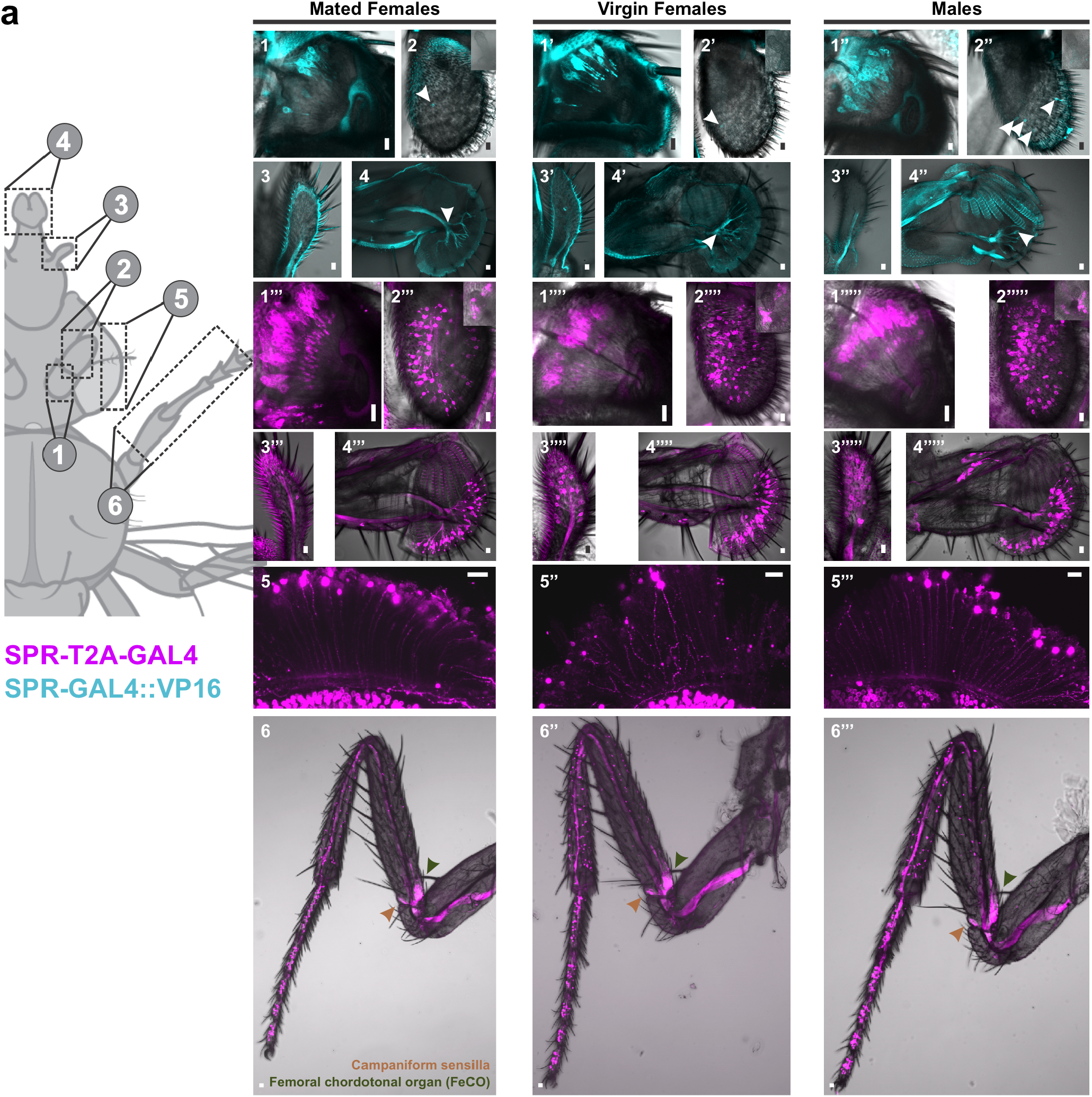
SPR-GAL4::VP16 and SPR-T2A-GAL4 expression throughout all primary sensory neurons. **(a)** Expression patterns of the bacterial artificial chromosome derived element SPR-GAL4::VP16 (cyan) and a CRISPR-Cas9 T2A-GAL4 insertion in the coding-intron of the sex peptide receptor (SPR-T2A-GAL4, magenta) in all sensory afferents in mated females, virgin females, and males. The left most diagram represents imaging plane for all images to the right, wherein: **1-1’’’’’** = auditory afferents; **2-2’’’’’** = olfactory, thermal, and hygrosensory afferents; **3-3’’’’’** = olfactory afferents; **4-4’’’’’** = gustatory afferents; **5-5’’’** = visual afferents; **6-6’’’** = proprioceptive and gustatory afferents. Driver expression in visual afferents and proprioceptive/gustatory afferents (in T1) were only tested for SPR-T2A-GAL4. Arrowhead(s) in **2-2’’’** and **4-4’’’** highlight the few neurons the express SPR-GAL4::VP16 in the 3^rd^-antennal segment (olfactory, thermal, and hygrosensory afferents), and the labellum (gustatory afferents), respectively. Neuron(s) that innervate the sacculus, a thermal/hygrosensory organ in the 3^rd^-antennal segment, are presented in the insets in the top right of **2-2’’’’’**. In all cases, scale bars = 10μm.

**Supplementary Fig. 5.**
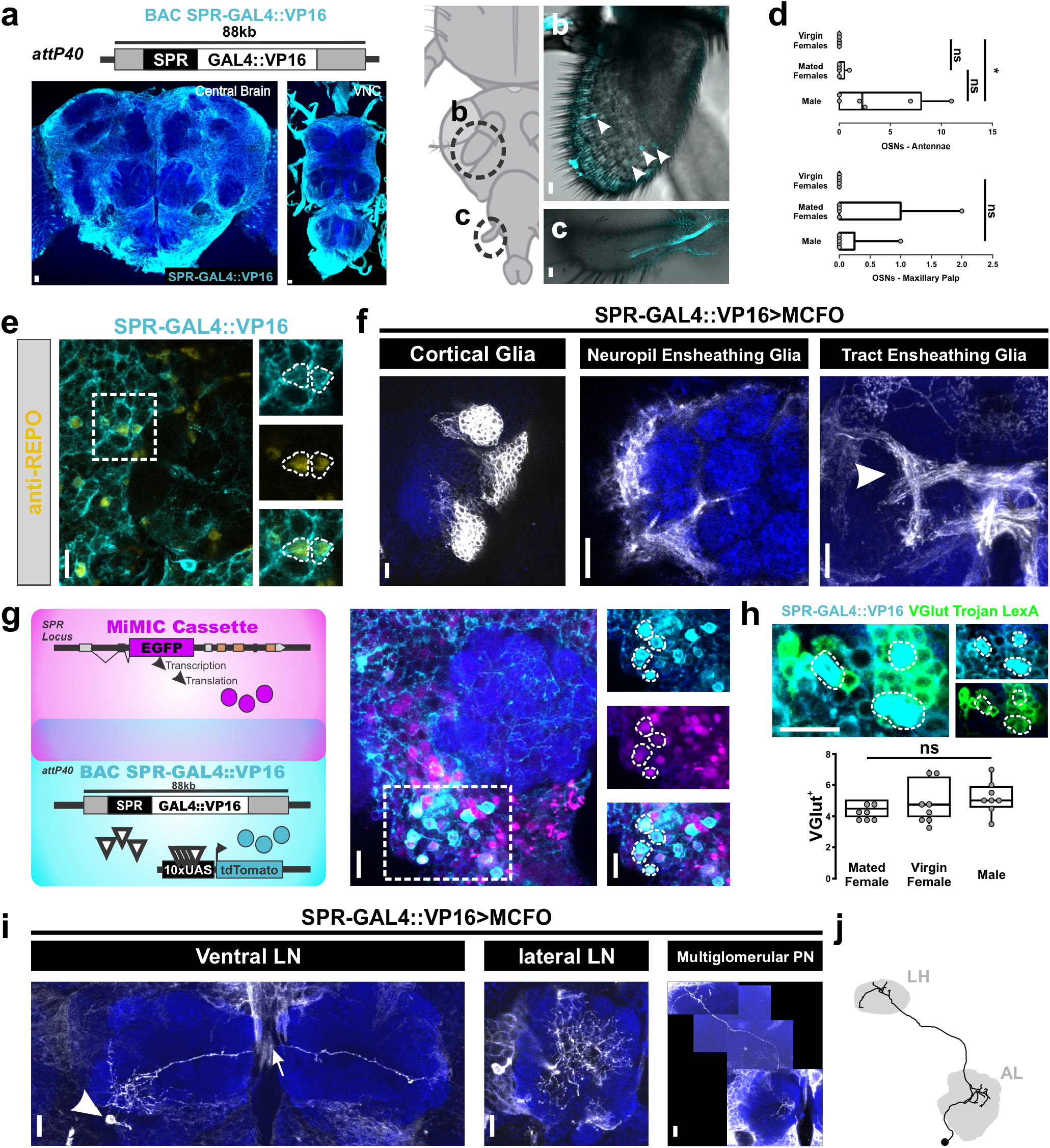
SPR-GAL4::VP16 expression throughout central brain circuitry with emphasis on AL expression. **(a)** Sex peptide receptor expression (SPR; cyan) as revealed using a bacterial artificial chromosome derived GAL4::VP16 element^90^. Note that this element contains the SPR locus and much of the surrounding genomic locus (~88kb total), and the GAL4::VP16 coding sequence was later inserted before the SPR stop site^90^. This element was then reintroduced at the attp40 landing site^90^. **(b-d)** SPR-GAL4::VP16 expression (cyan) in OSNs housed in the 3^rd^-antennal segment and maxillary palp. Female mating status does not affect the number of SPR-GAL4::VP16^+^ cells in antennae (p = 0.63; Holm-adjusted Dunn test), but males have significantly more SPR-GAL4::VP16^+^ cells in their antennae than virgin females (p = 0.05; Holm-adjusted Dunn test). However, the number of SPR-GAL4::VP16^+^ cells in the maxillary palp does not differ based on sex or mating status (p = 0.59; Kruskal-Wallis rank sum test). The discrepancy in the number of neurons of a given type observed between the SPR-T2A-GAL4 **(see Figure 7c & 7d)** versus the SPR-GAL4::VP16 drivers is likely a result of the non-native chromosomal topology, as well as potentially missing enhancer elements, of the SPR-GAL4::VP16 driver. **(e)** SPR-GAL4::VP16 (cyan) colocalizes with the general glial marker *reverse polarity* (anti-REPO; yellow). **(f)** SPR-GAL4::VP16 stochastic labeling highlights several glial subtypes, including cortical, neuropil ensheathing, and tract ensheathing glia. **(g)** Several ventral AL neurons are labeled through intersectional genetics experiments between an EGFP-insertion in the endogenous non-coding intron of SPR (MiMIC Cassette; magenta) and SPR-GAL4::VP16 (cyan). **(h)** At least a portion of the ventral AL neurons labeled by SPR-GAL4::VP16 are ventral glutamatergic LNs. The number of vesicular glutamate transporter-positive (VGlut^+^) SPR-GAL4::VP16 neurons does not statistically differ based on sex or mating status (p = 0.28, n = 8 (virgin females), 7 (mated females), and 8 (males); Kruskal-Wallis rank sum test). **(i)** SPR-GAL4::VP16 stochastic labeling confirms expression in ventral LNs, at least one lateral LN, and at least one ventral multiglomerular PN could be resolved. **(j)** Skeleton representation of the aforementioned SPR-GAL4::VP16 multiglomerular PN. In all cases: neuropil was delineated with anti-DN-cadherin staining; scale bars = 10μm.

**Supplementary Fig. 6.**
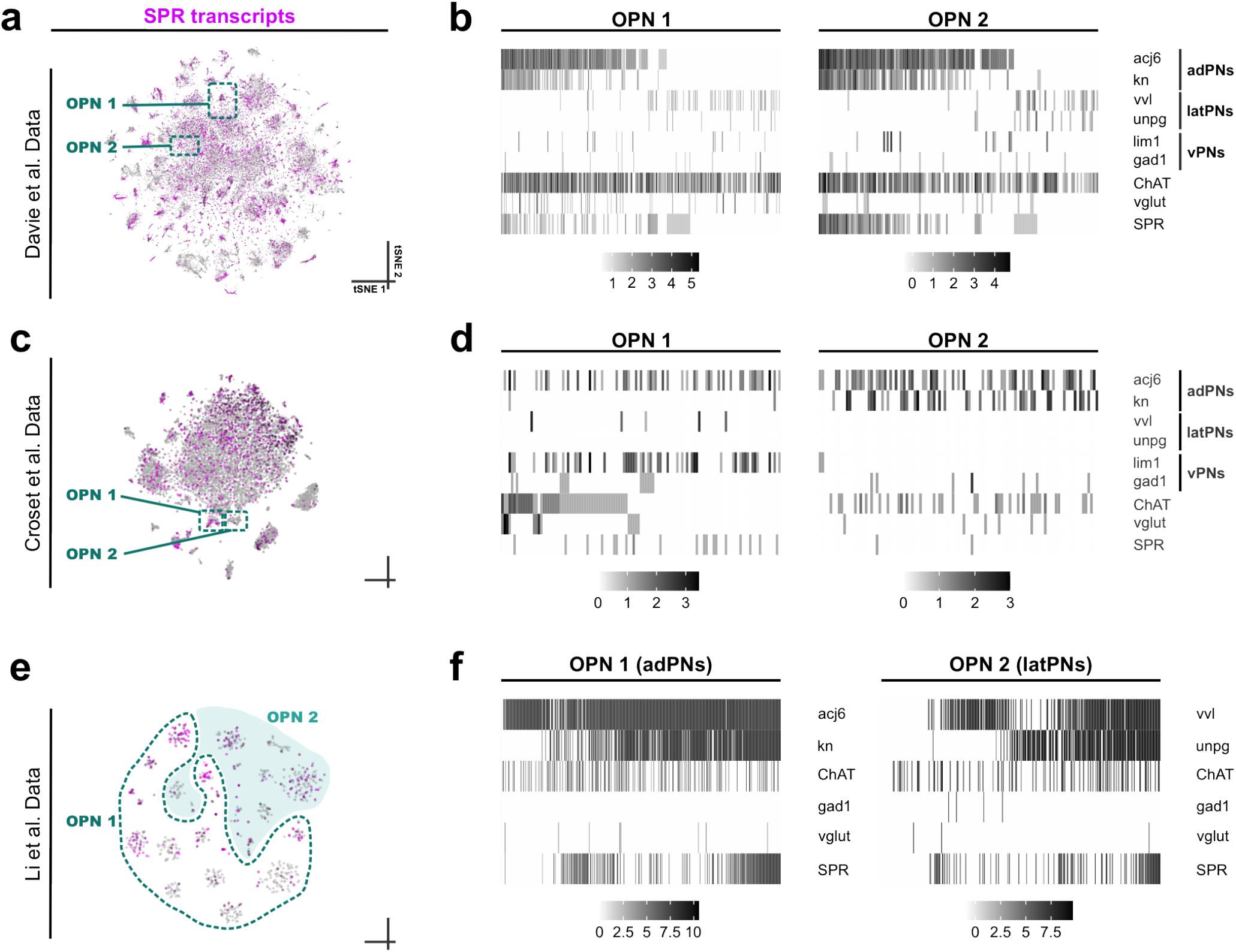
Sex peptide receptor (SPR) expression in independently generated projection neuron single-cell RNA-sequencing (scRNA-seq) datasets. **(a)** T-distributed stochastic neighbor embedding (tSNE) plot showing SPR expression (log-transformed and counts per million (CPM) normalized), wherein higher transcript levels are deeper magenta. **(b)** Heatmap showing transcript levels of anterodorsal projection neuron (adPN) marker genes *(acj6* and kn), lateral projection neuron (latPN) marker genes (*vvl* and *unpg*), ventral projection neuron (vPN) marker genes (*lim1* and *gad1*), choline acetyltransferase *(ChAT),* vesicular glutamate transporter (vglut), and SPR in olfactory projection neuron (OPN) clusters previously identified^93^. **(c)** As in **a**, visualization of SPR expression in Croset et al.^92^ scRNA-seq dataset. **(d)** As in **b**, heatmap showing transcript levels of various genes in OPN clusters previously identified^92^. **(e)** As in **a**, visualization of SPR expression in Li et al.^91^ OPN scRNA-seq data. **(f)** As in **b**, heatmap representation of transcript levels within adPN (OPN 1) and latPN (OPN 2) scRNA-seq clusters. Cluster boundaries were previously identified^91^. In all cases: Transcript levels are CPM normalized and log-transformed.

**Supplementary Fig. 7.**
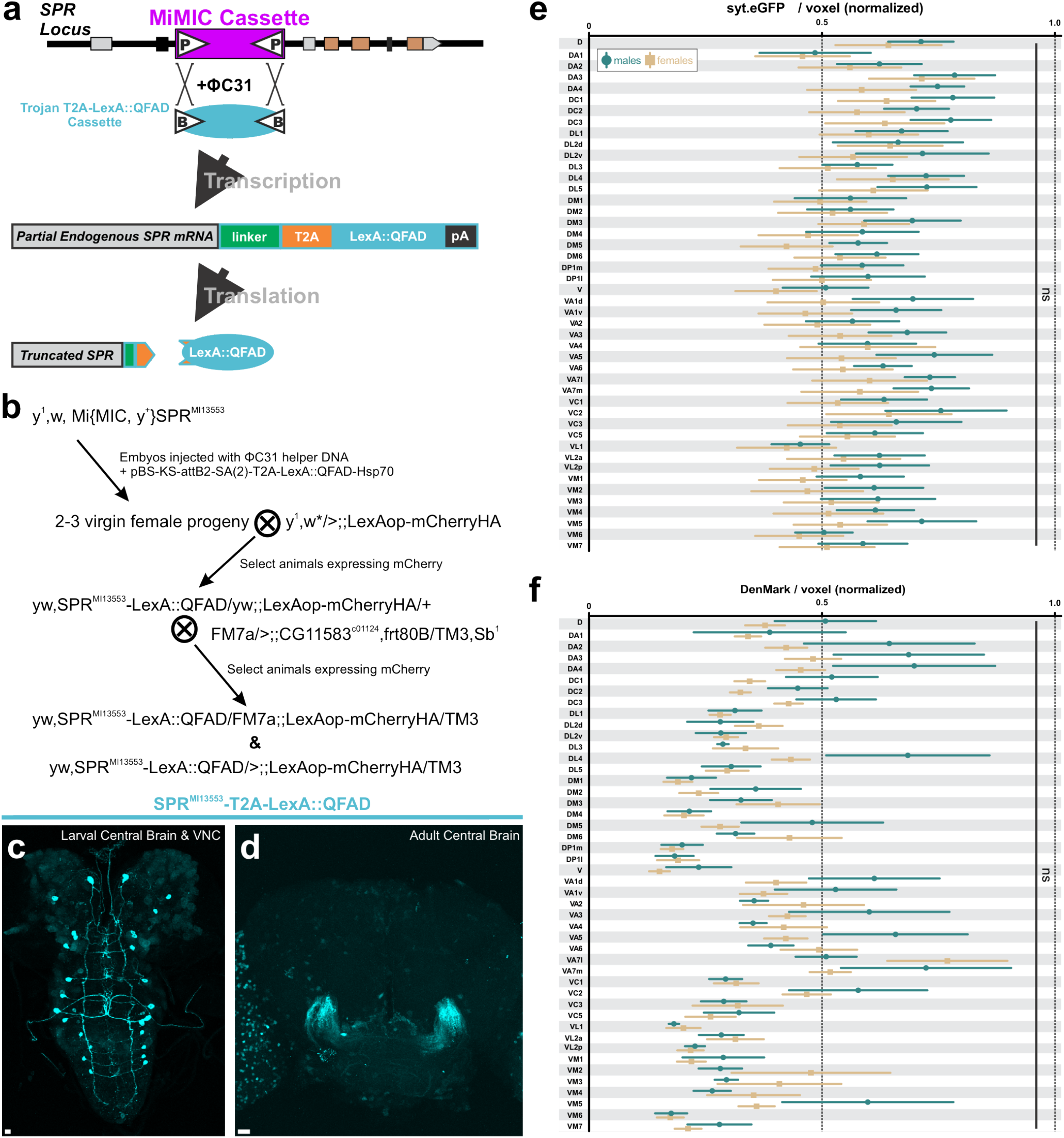
Generation of a sex peptide receptor (SPR) LexA::QFAD driver line via dual microinjection of a Trojan exon construct and ΦC31 recombinase. **(a)** Schematic representation of MiMIC cassette exchange for LexA::QFAD Trojan exon cassette, and subsequent LexA::QFAD expression in all cells that produce the sex peptide receptor (SPR). **(b)** Crossing scheme used to establish SPR^MI13553^-T2A-LexA::QFAD transgenics. **(c)** SPR^MI13553^-T2A-LexA::QFAD expression (cyan) in the larval central brain and ventral nerve cord. **(d)** SPR^MI13553^-T2A-LexA::QFAD expression (cyan) in the adult central brain. Note that while soma labeled by this driver are faintly reliably resolvable, neural processes are generally unresolvable in the adult. **(e & f)** R32F10-GAL4 synaptotagmin-eGFP (syt.eGFP) and DenMark voxel density do not display sexual dimorphism across glomeruli (syt.eGFP: p = 0.0634; DenMark: p = 0.4347; n = 3 brains, 6 ALs (male), 4 brains, 8 ALs (females); two-way ANOVA). In all cases, scale bars = 10μm.

